# Kinetic and Redox Characterization of KRAS G12C Inhibition

**DOI:** 10.1101/2022.04.03.486828

**Authors:** Minh V. Huynh, Derek Parsonage, Tom E. Forshaw, Venkata R. Chirasani, G. Aaron Hobbs, Hanzhi Wu, Jingyun Lee, Cristina M. Furdui, Leslie B. Poole, Sharon L. Campbell

**Affiliations:** Department of Biochemistry & Biophysics, University of North Carolina at Chapel Hill, Chapel Hill, NC 27599, USA; Department of Biochemistry, Wake Forest School of Medicine, Winston-Salem, NC 27157, USA; Department of Internal Medicine, Section on Molecular Medicine, Wake Forest School of Medicine, Winston-Salem, NC 27157, USA; Department of Pharmacology, University of North Carolina at Chapel Hill, Chapel Hill, NC 27599, USA; Lineberger Comprehensive Cancer Center, University of North Carolina at Chapel Hill, Chapel Hill, NC 27599, USA; Wake Forest Baptist Comprehensive Cancer Center, Winston-Salem, NC 27157; Center for Redox Biology and Medicine, Wake Forest School of Medicine, Winston-Salem, NC 27157, USA

**Author notes:** Medical University of South Carolina, Charleston, SC 29425, USA.

**Keywords:** KRAS^G12C^, oxidation, covalent inhibitors, lung cancer, thiol alkylators, RAS GTPase

## Abstract

The development of mutant-selective inhibitors for the *KRAS^G12C^* allele has generated considerable excitement. These KRAS^G12C^ inhibitors covalently engage the mutant C12 thiol located within the phosphoryl binding loop of RAS, locking the KRAS^G12C^ protein in an inactive state. While clinical trials of these inhibitors have been promising, mechanistic questions regarding the reactivity of this thiol remain, motivating the present studies. Measurement of the C12 thiol pK_a_ by NMR and an independent biochemical assay found a depressed pK_a_ (relative to free cysteine) of 7.6 consistent with its susceptibility to chemical ligation. Using a novel and validated fluorescent KRAS^Y137W^ variant amenable to stopped-flow spectroscopy, we characterized the kinetics of KRAS^G12C^ fluorescence changes upon addition of ARS-853 or AMG 510, noting that ARS-853 addition at 5°C elicited both a rapid first phase (attributed to binding, yielding a K_d_ of 36.0 ± 0.7 μM), and a second, slower pH-dependent phase taken to represent covalent ligation. Consistent with the lower pK_a_ of the C12 thiol, we found that reversible and irreversible oxidation of KRAS^G12C^ occurred readily both *in vitro* and in the cellular environment, preventing the covalent binding of ARS-853. Moreover, we found that oxidation of the KRAS^G12C^ thiol to sulfinic acid alters RAS conformation and dynamics to be more similar to KRAS^G12D^ in comparison to the unmodified protein, as assessed by molecular dynamics simulations. Taken together, these findings provide insight for future KRAS^G12C^ drug discovery efforts as well as identifying the occurrence of G12C oxidation with currently unknown biological ramifications.

## INTRODUCTION

Mutations in *KRAS* are known to be key drivers in some of the most lethal human cancers. As such, the development of potent anti-RAS therapies has been an exciting prospect for more than 30 years (1–3). Recently, anti-RAS drug discovery efforts have taken a massive stride forward with the development of small molecule inhibitors that selectively target tumors carrying the *KRAS^G12C^* mutant allele (4,5). The *KRAS^G12C^* mutation accounts for approximately 12% of *KRAS* mutations across all human cancers and is the predominant mutation in *KRAS*-mutant non-small cell lung adenocarcinomas (NSCLC). These novel RAS inhibitors take advantage of the mutated cysteine residue as a covalent tether and bind to KRAS in a previously unknown pocket now known as the Switch II pocket (6,7). With these compounds effectively locking KRAS^G12C^ in an inactive GDP-bound state, several of these covalent inhibitors have achieved exciting clinical success, with AMG 510 and MRTX849 both showing potent anti-tumor activity as single agents and in combination studies (8,9). Excitingly, AMG 510 (Lumakras™, Amgen, Inc) has received accelerated approval by Food and Drug Administration in 2021 for treatment of locally advanced or metastatic NSCLC carrying the *KRAS^G12C^* mutation.

However, despite the potent activity of these clinical compounds, the kinetics and reactivity of KRAS^G12C^ inhibitors in general is poorly understood. Previous studies assessing the reactivity for this class of KRAS inhibitors were performed using the inhibitor ARS-853 but faced challenges such as limited inhibitor solubility and lack of intrinsically fluorescent tryptophan residues which impeded analyses (10). Additionally, the exact mechanisms behind the observed reactivity of KRAS^G12C^ to alkylating inhibitors such as ARS-853 remain unclear (11). As the understanding of these processes is critical for the development of future KRAS inhibitors, we sought to resolve these gaps in knowledge. In this study, we characterized the pK_a_ and reactivity of the mutant G12C residue via several direct and indirect methods and demonstrate that the KRAS^G12C^ thiol exhibits a depressed pK_a_. We established and validated a novel Y137W variant of KRAS^G12C^ that allows for direct evaluation of KRAS^G12C^ inhibitor binding and ligation kinetics. Further, we found that oxidation of the G12C cysteine occurs in solution and in cell cultures and blocks chemical ligation by covalent KRAS^G12C^ inhibitors. Lastly, we found that oxidation of KRAS^G12C^ to the sulfinic acid state mimics KRAS^G12D^ by molecular dynamics (MD) analyses. Together, our study establishes an assay for future kinetic assessment and development of KRAS inhibitors and suggests a role for oxidation in the regulation of KRAS^G12C^-specific biological functions.

## RESULTS

### KRAS^G12C^ exhibits a perturbed cysteine pK_a_ by chemical modification and NMR

Previous studies from our group and others have shown that several cysteine residues in a subset of RAS superfamily GTPases have a lower pK_a_ relative to that of a free cysteine (12–15). With a lowered pK_a_, these cysteines are more populated in the reactive thiolate state at physiological pH. We hypothesized that the mutated G12C residue in KRAS would also display a perturbed pK_a_ owing in part to its position within the RAS nucleotide binding site. To test this hypothesis, we assessed the cysteine reactivity of the G12C residue by monitoring the reaction of KRAS with the fluorogenic agent 4-aminosulfonyl-7-fluoro-2,1,3-benzoxadiazole (ABD-F). This electrophile reacts with the thiolate state of cysteine residues approximately 10-fold faster than with the protonated state. The core guanine nucleotide binding domain of native KRAS contains three cysteine residues (C51, C80, C118). While C51 and C80 are buried, C118 is solvent accessible (12). To eliminate the possibility of ABD-F modification at the solvent accessible site, we introduced a C118S mutation. We previously have shown that this mutation does not impact KRAS biochemical function (16). We tested ABD-F reactivity across a pH range (5.8 to 8.2) and found that KRAS^G12C^ is susceptible to ABD-F modification in a pH-dependent manner (**Fig. 1A**). From fitting the initial rates of ABD-F modification at different pH values, we estimated the pK_a_ for the KRAS^G12C^ thiol to be 7.6 ± 0.1 (**Fig. 1B**). The negative control KRAS^G12S/C118S^ did not show appreciable reactivity to ABD-F.

**Figure 1.**
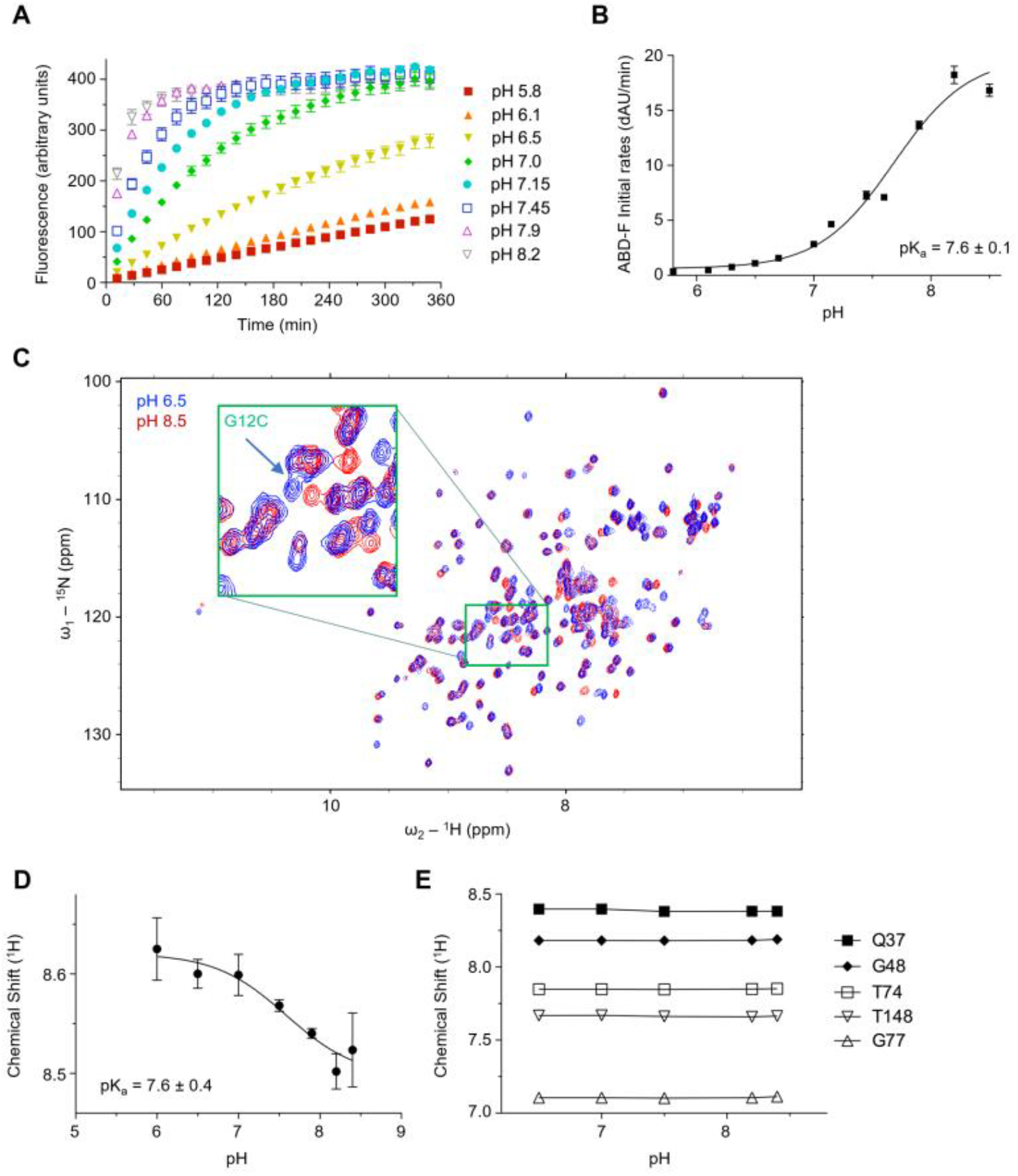
KRAS^G12C^ exhibits a depressed pK_a_ relative to free cysteine. **(A)** Reaction of ABD-F with KRAS^G12C/C118S^ over a pH range from 5.5 to 8.5 (KRAS at 10 μM, ABD-F at 1 mM). Data were collected over 6 h in triplicate with absolute fluorescence values plotted. Data shown are representative from three or more independent experiments. **(B)** Reaction rates of modification of KRAS^G12C/C118S^ by ABD-F plotted against pH. The variation in initial rates of ABD-F modification across pH was analyzed using GraphPad Prism, yielding a *pK_a_* value of 7.6 ± 0.1 for the KRAS^G12C^ thiol. Data shown are averaged from three or more independent experiments. Error bars, mean ± s.e.m. **(C)** ^1^H-^15^N 2D NMR HSQC overlay of ^15^N-labeled KRAS^G12C/C118S^ in the GDP-bound state at pH 6.5 (blue) vs. pH 8.5 (red). The green insert indicates the location of the KRAS^G12C^ amide cross-peak. **(D)** NMR pK_a_ determination of the KRAS^G12C^ thiol from ^1^H-^15^N amide proton chemical shift perturbations. The p*K*_a_ of the KRAS^G12C^ thiol was estimated to be 7.6 ± 0.4. Data shown are averaged from three or more independent experiments. Error bars, mean ± s.e.m. **(E)** ^1^H-^15^N amide proton chemical shift perturbations of control residues across the pH titration range used to determine the KRAS^G12C^ thiol pK_a_. Data shown are representative from three or more independent experiments.

While the reactivity of a cysteine thiol is partially dependent on its surrounding electrochemical environment, the electrophilicity and binding orientation of reacting compounds such as ABD-F will also influence the rate of cysteine modification (17). Hence, we employed an independent, NMR-based approach to directly assess the pK_a_ of C12 in the absence of small molecules. With NMR backbone assignments readily available for KRAS^G12C^ (18,19), we acquired 2D NMR ^1^H-^15^N Heteronuclear Single Quantum Coherence (HSQC) spectra and were able to monitor chemical shift perturbations for the C12 residue across a similar pH range as used for ABD-F modification (**Fig. 1C**). Chemical shifts for the mutant KRAS^G12C^ C12 NH cross peaks demonstrated responsive changes across the pH range tested; negligible shifts observed for several control residues indicated minimal structural alterations to KRAS (**Fig. 1D, E**). From these data, we found that direct monitoring of KRAS^G12C^ chemical shifts by NMR yielded a pK_a_ of 7.6 ± 0.4, similar to the value obtained from ABD-F modification.

### Design and validation of a fluorescent form of KRAS^G12C^

Understanding the mechanism behind the observed reactivity of the cancer-associated G12C residue is highly relevant for current and future KRAS^G12C^-specific inhibitor design. However, our understanding of the kinetics of binding and ligation by these KRAS^G12C^-specific inhibitors is currently quite limited. Given the lack of any naturally occurring tryptophan residues in KRAS which could serve as fluorescent sensors of inhibitor association amenable to kinetic analysis, we set out to identify possible sites for introducing a tryptophan as an intrinsic probe in KRAS. Analyzing the sequence conservation of several other RAS GTPases, we found that several members of the RAS subfamily contain a tryptophan at the position analogous to KRAS Y137 (**Fig. 2A**). Of note, Y137 in KRAS is located at the end of helix α4 and is not involved in any intramolecular interactions. Analysis of published crystal structures for several of these RAS subfamily members show that the analogous tryptophan residues occupy a similar side-chain position, suggesting that mutation of Y137 in KRAS to tryptophan is a conservative substitution that will not likely affect RAS structure or function (**Fig. 2B, C**). Additionally, Y137 is distal to the RAS nucleotide binding site and thus is unlikely to interfere with kinetic studies interrogating the cysteine reactivity of KRAS^G12C^.

**Figure 2.**
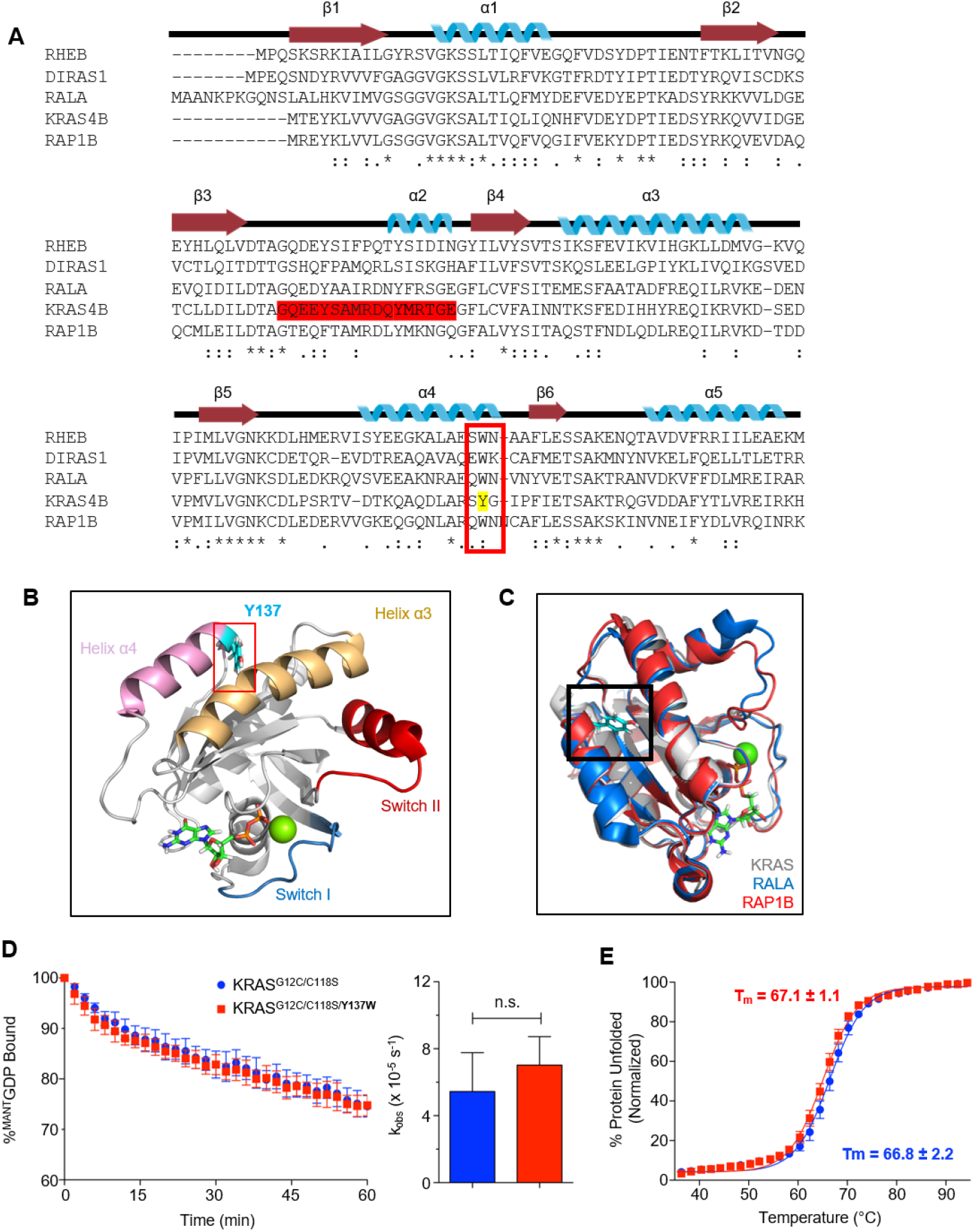
Characterization of the Y137W substitution in the C118S variant of KRAS^G12C^. **(A)** Sequence alignment of Ras subfamily GTPases showing conservation in the region of KRAS Y137. Alignment shows key alpha-helix and beta-sheet structural motifs (KRAS residues 1-165). The red box highlights the location of Y137 in KRAS and the corresponding tryptophan residue present in related family members. **(B)** 3D location of residue Y137 in KRAS (PDB: 4LPK). Y137 is shown in sticks and highlighted in cyan, bound GDP is shown in sticks, and a bound calcium ion is shown in green. Switch I: teal, Switch II: red. Ribbon diagram was rendered using PyMOL. **(C)** Structural overlay of GDP-bound KRAS4B (PDB: 4LPK), RAL A (PDB: 6P0J), and RAP1B (PDB 3X1W) highlighting location of the residues equivalent to KRAS Y137 (black box). **(D)** Representative intrinsic nucleotide dissociation monitoring the decrease in ^mant^GDP fluorescence over time after mixing with non-fluorescent GDP, comparing KRAS^G12C/C118S^ with and without the Y137W mutation, shows minimal effects of the Y137W mutation on nucleotide exchange activity. Data shown are representative from three or more independent experiments. Error bars, mean ± s.e.m. **(E)** Comparison of protein stability by circular dichroism thermal denaturation for KRAS^G12C/C118S^ vs. KRAS^G12C/C118S/Y137W^. The similar midpoint Tm values indicate that Y137W does not notably impact protein thermal stability. Data shown are representative from three or more independent experiments. Error bars, mean ± s.e.m.

To verify that this mutation does not significantly alter protein function, we performed nucleotide exchange analysis on KRAS^G12C^ with and without the Y137W mutation. We found that the GDP nucleotide dissociation rates for KRAS^G12C/Y137W^ were similar to the control KRAS^G12C^ (**Fig. 2D**). We also tested the effects of the mutation on the thermal stability of KRAS^G12C^ using circular dichroism (CD) spectroscopy and did not observe significant changes in the protein melting temperature or cooperativity of the unfolding transition (**Fig. 2E**). Further, MD analyses indicate that introduction of the Y137W substitution into KRAS^G12C^ does not alter KRAS structure or dynamics (**Fig. S1**). Together, these data indicate that the Y137W substitution does not significantly alter RAS nucleotide binding or stability, supporting the use of this fluorescent mutant to analyze kinetic behavior.

### Stopped-flow kinetic studies of KRAS^G12C^ interaction with ARS-853 show both fast and slow processes and a functional pK_a_ for inhibitor ligation

Generation of the Y137W mutation in KRAS^G12C^ enabled use of this fluorescent residue as a tool to directly monitor fluorescence changes associated with inhibitor binding and ligation to KRAS^G12C^, including the KRAS^G12C^-specific inhibitor ARS-853. ARS-853 is an early model compound that is similar to clinically relevant KRAS^G12C^-specific inhibitors with reactive acrylamide moieties and which has undergone previous kinetic assessment (7,10). Further, we introduced mutations at the other three cysteine residues of KRAS^G12C^ (C51S/C80L/C118S) to generate a “Cys-light” protein to eliminate off-target inhibitor ligation in this assay (5), generating a KRAS^G12C/C51S/C80L/C118S/Y137W^ variant (referred to as KRAS^CCLW^). The “Cys-light” version of KRAS^G12C^ was previously shown to have only minimal effects on overall protein structure (5). The protein was freshly reduced using 1,4-dithiothreitol (DTT), followed by removal of the DTT prior to kinetic analyses.

Stopped flow kinetic experiments conducted at 5°C to capture data for the fastest reactions revealed that addition of ARS-853 to pre-reduced KRAS^CCLW^ resulted in very rapid, concentration-dependent fluorescence changes up to ~250 ms (**Fig. 3A** and **3B**). We confirmed that the protein and not the inhibitor was responsible for the observed fluorescence changes; the kinetic traces of all reaction mixes, when extrapolated back to the time of mixing, yielded the same fluorescence as KRAS mixed with buffer. No pH dependence of reaction rates was observed, beyond modestly lower rates at pH 9 (**Fig. 3B**).

**Figure 3.**
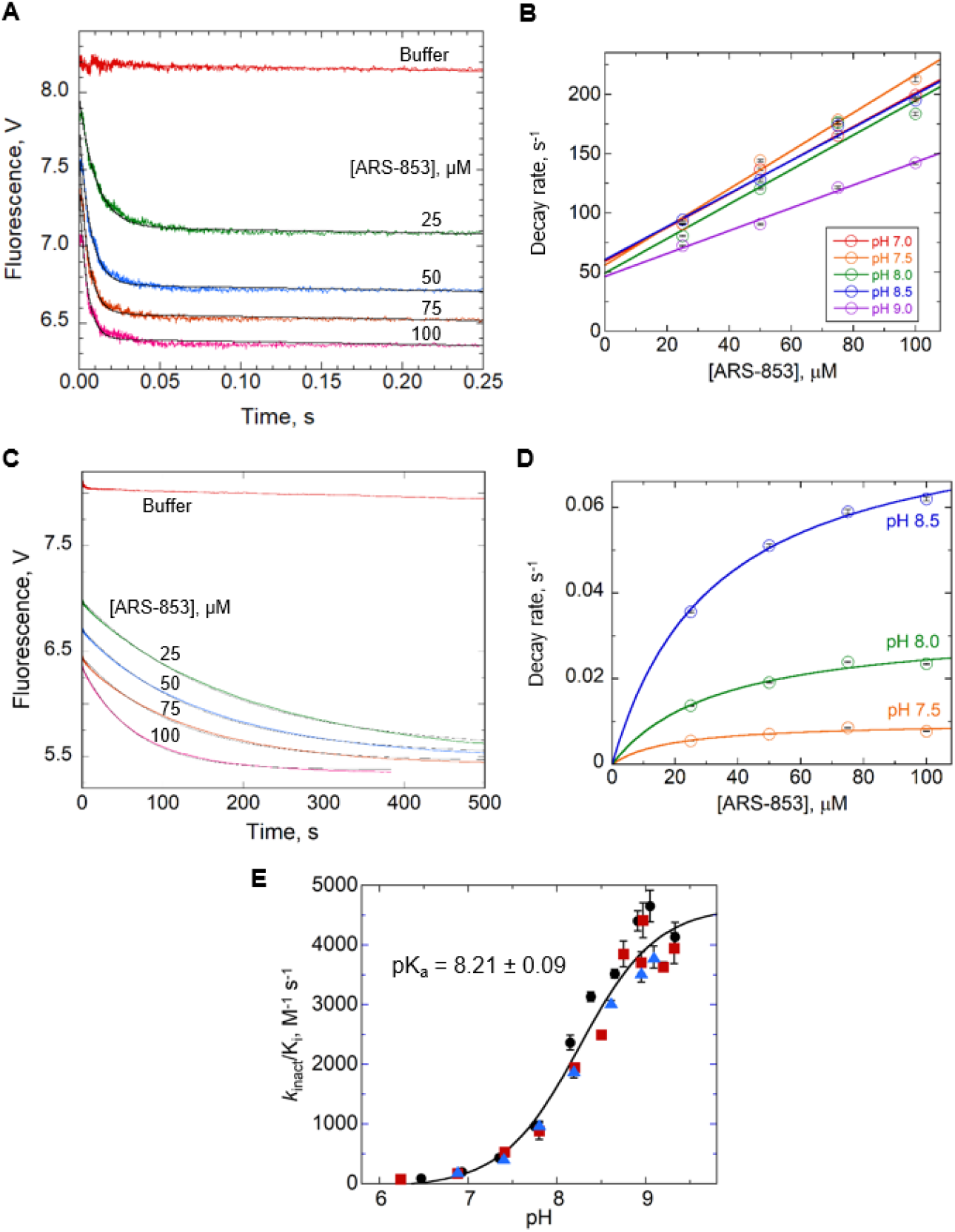
Stopped-flow kinetic studies of KRAS^CCLW^ fluorescence changes upon ARS-853 addition show a very fast fluorescence decrease followed by a slower decrease, and a functional pK_a_ of ~8.2 for the slow step. **(A)** Stopped-flow kinetic data of 5 μM KRAS^CCLW^ (G12C,”Cys-light”, Y137W) mixed with buffer or ARS-853 at 25, 50, 75 and 100 μM at 5°C and pH 7.5 demonstrate an initial fast drop in fluorescence (< 50 ms). Data shown are representative of three or more independent experiments. **(B)** Plots of the pseudo first order rate constant *versus* ARS-853 concentration at five pH values (average plus or minus the standard error of each) yield a linear fit and an absence of a pH effect up to pH 8.5. **(C)** A further decrease in fluorescence beyond that seen in A occurs over ~500 s at 5°C (same colors and concentrations as in **A**). Data shown are representative of three or more independent experiments. **(D)** Pseudo-first order rate constants at 5°C fit to a hyperbola are consistent with a saturable step (interpreted as covalent modification). As shown, rate constants are strongly affected by pH. **(E)** Second-order rate constants (k_inact_/K_i_) for the slow step of KRAS^CCLW^ and ARS-853 reactions were fit to a Boltzmann sigmoidal; the ARS-853 functional pK_a_ was found to be 8.21 ± 0.09 at 20°C. Three colors and marker shapes represent three independent trials, and each data point reports the mean ± standard error of the mean for the k_inact_/K_i_ value derived from multiple [ARS-853]; all data were used in the final fit.

Under these conditions, a second, much slower decrease in fluorescence is also observed (from ~1-500 s, **Fig. 3C**); in this case, data collected at various pH values show a strong pH effect (**Fig. 3D**). For this slower step, which can be observed at both 5°C and 20°C, the first order decay rate plotted *versus* inhibitor concentration indicates a saturable interaction (**Fig. 3D**). Kinetic profiles of covalent enzyme inhibition typically yield a hyperbolic dependence on inhibitor concentration (Equation 1).

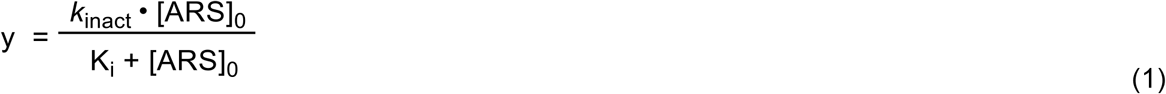

In this equation, y is the first order rate of decay derived from exponential fits of the data, *k*_inact_ is the rate constant of the irreversible chemical step, K_i_ is the equilibrium constant corresponding to reversible binding, and [ARS]_o_ is the initial concentration of the ARS-853 inhibitor (11). Plotting *k*_inact_/K_i_ (which can be determined even when K_i_ is very high) *versus* pH indicates a “functional” pK_a_ (associated with the ligation reaction) at 20°C of 8.21 +/- 0.09 (**Fig. 3E**). A very similar value of 8.15 ± 0.13 is obtained at 5°C (**Fig. S2**). Note that these functional pK_a_ values are 0.6-0.7 pH units higher than the C12 pK_a_ of the of the inhibitor-free enzyme determined by NMR.

The simplest interpretation of the fast reaction with linear dependence on [I] (observable at 5°C with ARS-853, **Fig. 3A** and **3B**) is that these fluorescence changes represent reversible binding of the inhibitor to the protein. The much slower reaction, with hyperbolic dependence on [I] (**Fig. 3C** and **3D**), represents the irreversible chemical reaction between the two (Equation 2).

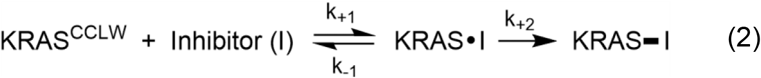

Note that in this kinetic model, k_+1_ represents k_on_ and k_-1_ represents k_off_ for reversible inhibitor binding, and K_d_ can be calculated from the ratio of the two. In addition, k_+2_ represents k_inact_ and k_-2_, not shown in the model, is 0 due to the irreversible formation of the covalent bond between KRAS and inhibitor. To establish the most accurate kinetic parameters from these data collected at 5°C, we conducted global analyses of the raw data (including both the fast and slow reaction data and all inhibitor concentrations) using KinTek Explorer and the kinetic model of Equation 2. Individual rate constants and the computed K_d_ are shown in Table 1.

**Table 1.** Rate constant values from global fits of data shown in **Fig. 3A** using KinTek Explorer.

| pH | $k_{+1}$ | $k_{-1}$ | $k_{+2}$ | $K_d$ |
| --- | --- | --- | --- | --- |
| 6.98 | $1.35 \pm 0.01$ | $49.7 \pm 0.3$ | $0.0073 \pm 0.0004$ | $36.8 \pm 0.4$ |
| 7.45 | $1.69 \pm 0.01$ | $51.1 \pm 0.3$ | $0.0169 \pm 0.0005$ | $30.2 \pm 0.3$ |
| 7.97 | $1.50 \pm 0.01$ | $51.2 \pm 0.3$ | $0.038 \pm 0.006$ | $34.2 \pm 0.3$ |
| 8.46 | $1.45 \pm 0.01$ | $61.7 \pm 0.4$ | $0.099 \pm 0.001$ | $42.6 \pm 0.4$ |
| 8.95 | $0.976 \pm 0.009$ | $75.1 \pm 0.4$ | $0.280 \pm 0.003$ | $77.0 \pm 0.8$ |

Averaging the K_d_ values determined from pH 7.0 to 8.5 yields a K_d_ of 36.0 ± 0.7 μM at 5°C. Given the kinetic model used (Equation 2), this is also the K_i_ value for ARS-853 at 5°C. At 20°C, we were no longer able to track the fast rate and therefore could not determine the K_d_ directly. Instead, we determined K_i_ and k_inact_ values by conducting hyperbolic fits as recommended by Johnson (20), and then used the k_inact_/K_i_ values in the determination of the functional pK_a_ (**Fig. 3E** and **Fig. S2**).

As several KRAS^G12C^-targeting inhibitors are now in late-stage clinical trials (8,9,21), we next assessed the more clinically relevant inhibitor AMG 510 by rapid reaction kinetics for comparison with ARS-853. We again observed a rapid drop in fluorescence upon mixing AMG 510 with the protein. However, this fluorescence decrease, unlike for ARS-853, occurred within the deadtime of the instrument (~1.5 ms) at 5°C, indicating that binding equilibration is much faster with this compound (**Fig. 4A**). Evaluating the kinetics at 20°C, the rates of fluorescence decay were higher at every concentration with AMG 510 relative to ARS-853, although the K_i_ was also higher for AMG 510 than for ARS-853 (**Fig. 4B** and **Fig. S3**). Evaluation of the *k*_inact_/K_i_ values for AMG 510 *versus* ARS-853 at three different pH values showed that AMG 510 exhibits a >5-fold larger value than ARS-853 independent of pH (**Table S1**, **Fig. S3**).

**Figure 4.**
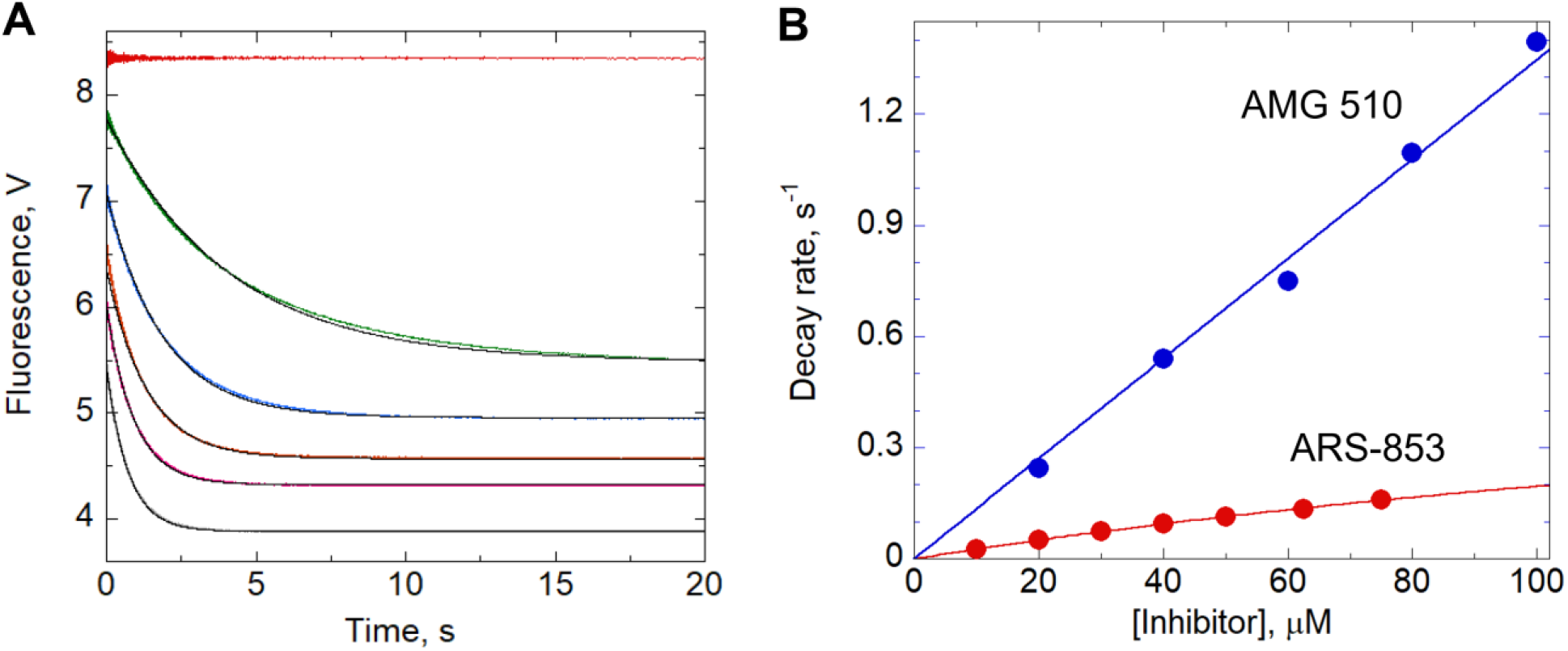
Kinetic studies of KRAS^CCLW^ fluorescence changes with increasing inhibitor concentrations (at 20°C) are faster for AMG 510 than for ARS-853. **(A)** Stopped-flow kinetic data of 5 μM KRAS^CCLW^ mixed with buffer or AMG 510 at 20, 40, 60, 80 and 100 μM at 20°C and pH 8.61 (faster fluorescence decreases reflect higher concentrations of inhibitor) demonstrate faster kinetics for AMG 510 reactions than for “slow step” reactions with ARS-853. **(B)** Plots of the observed rates *versus* inhibitor concentrations suggest very high K_i_ values for both inhibitors (K_i_ > 100 μM) under these conditions. Fitted values for k_inact_/K_i_ are 2770 ± 10 and 14000 ± 100 M^-1^ s^-1^ with ARS-853 and AMG 510, respectively.

In order to evaluate the reaction kinetics for comparison with a previous study in which K_i_ and k_inact_ were determined using a very low concentration of ARS-853 (8 μM) and varying high concentrations of protein (Hansen *et al*. (10)), we conducted experiments at 20°C (the Hansen study was conducted at room temperature) and pH 7.5 using the reaction buffer described in their study (20 mM HEPES, with 150 mM NaCl, 1 mM DTT and 1 mM MgCl_2_). The buffer used in our studies (MHT buffer in Materials and Methods), lacks DTT and has 50 mM NaCl and 5 mM MgCl_2_, also included 50 mM each of MES and Tris in addition to 50 mM HEPES to allow experiments at varying pH. Surprisingly, while the two buffers yielded overlapping results at ARS-853 concentrations up to 150 μM, the DTT-containing buffer exhibited visible saturation behavior at higher ARS-853 concentrations of 200 μM and above whereas the rate constant in MHT buffer continues to rise nearly linearly up to 300 μM (**Fig. S4**). Comparing our results with those from Hansen *et al*., we found that, in the DTT-containing buffer, our data yielded a value for K_i_ of 142 ± 19 μM) compared to the previous paper (200 ± 90 μM (10)) and an approximately two-fold higher *k*_inact_/K_i_ (510 ± 75 rather than 250 M^-1^ s^-1^, **Table S2**). Our experiments reveal how dependent on the buffer conditions these values are, as the K_i_, along with the *k*_inact_, appears to be much higher in our MHT buffer at pH 7.5 (on the other hand, *k*_inact_/K_i_ in MHT buffer is only modestly different, at 336 ± 45 M^-1^ s’^1^).

### *KRAS^G12C^ is sensitive to oxidative modification* in vitro *and in NIH 3T3 cells*

KRAS and other RAS superfamily GTPases have previously been shown to be sensitive to cysteine oxidation both *in vitro* and in cellular contexts (12–15). With the observed sensitivity of KRAS^G12C^ to chemical ligation by acrylamides like AMG 510 and ARS-853 and our NMR data demonstrating the lowered pK_a_ of the G12C cysteine, we hypothesized that this cysteine could also be sensitive to modification by intracellular oxidizing agents. Further, as current KRAS^G12C^ inhibitors are electrophilic and designed to specifically alkylate the free, nucleophilic G12C thiol for inhibition, we hypothesized that oxidation of this residue would block covalent modification.

To test this, we utilized electrospray ionization time-of-flight (ESI-TOF) mass spectrometry to detect specific modifications at the C12 residue by ARS-853 in the presence and absence of H_2_O_2_-mediated oxidation. Using *in vitro* studies with recombinant KRAS^CCLW^, we found that the reduced form of the protein reacted rapidly with a 1.5-fold excess of ARS-853 and formed an adduct within 5 min (**Fig. 5A**). In addition, the reduced protein (-SH) was oxidized by reaction with H_2_O_2_ (at 1:1 or up to 10-fold excess) within 10 min to generate a mixture of sulfenic (-SOH) and sulfinic (-SO_2_H) acid modifications (**Fig. 5A**). Data obtained at intermediate concentrations confirm the sensitivity of the G12C cysteine residue to oxidation and hyperoxidation (**Fig. S5**). The oxidized KRAS^CCLW^ -SOH, which displays both nucleophilic and electrophilic properties, reacted rapidly with the selective sulfenic acid-reacting molecule dimedone but did not react with ARS-853 (**Fig. 5A**). Additionally, and as expected, KRAS^CCLW^ -SO_2_H did not react with ARS-853. Taken together, our findings indicate that H_2_O_2_-oxidized KRAS^CCLW^ likely does not react with other alkylating KRAS^G12C^-specific inhibitors either given similarity of the warheads.

**Figure 5.**
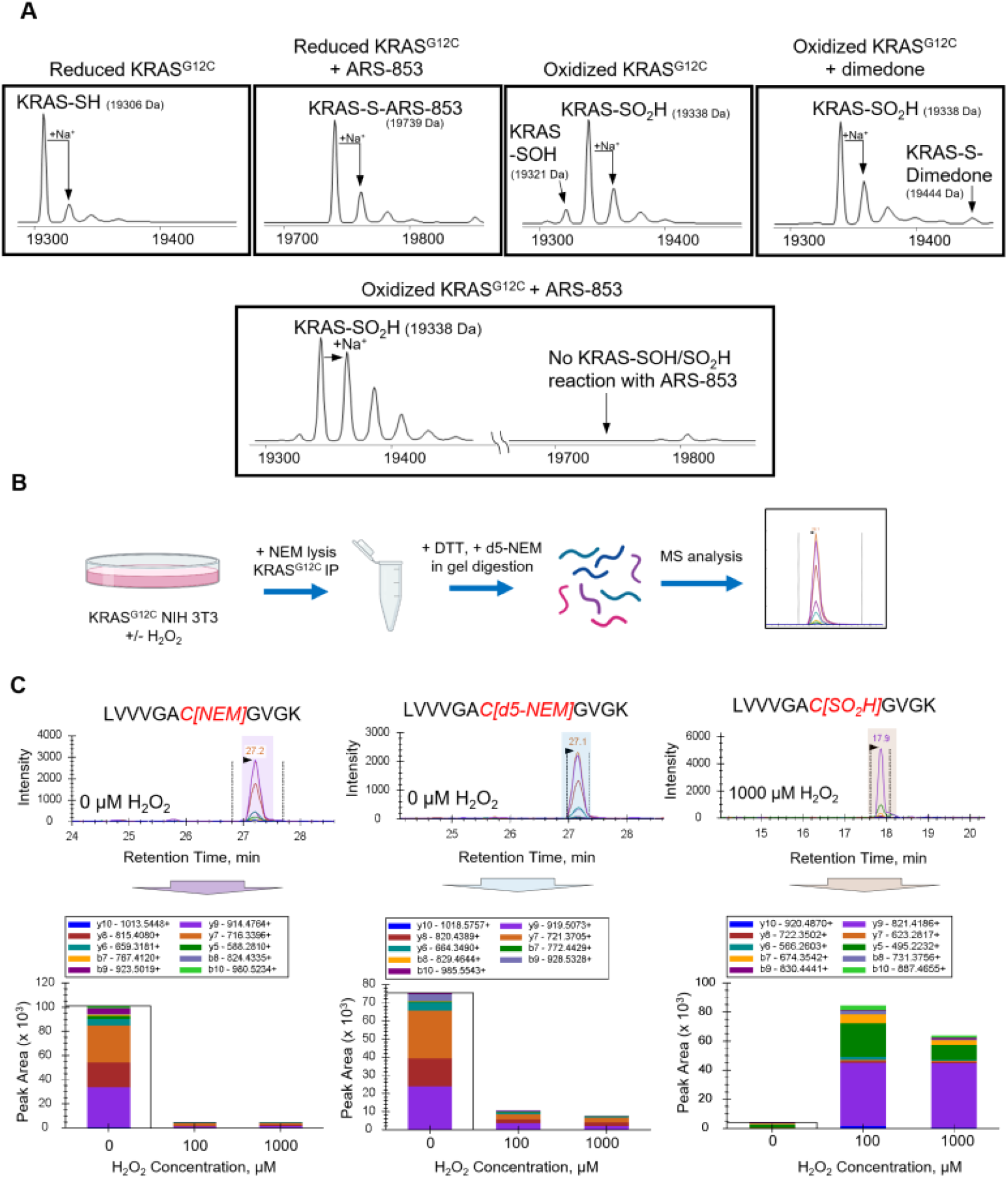
Inhibitor reactivity and redox state of KRAS^G12C^ *in vitro* and in NIH 3T3 cells. **(A)** As shown by mass spectrometry, pre-reduced KRAS^CCLW^ (as used in Fig. 3) reacts rapidly with ARS-853 (second panel, incubated 5 min with a 10-fold excess of the inhibitor). Oxidation of KRAS^CCLW^ by reaction with H_2_O_2_ (50-fold excess of H_2_O_2_ added for 10 min) produced a mixture of sulfenic acid (-SOH; confirmed by reaction with dimedone) and sulfinic acid (SO_2_H) forms, which did not react with ARS-853. **(B)** Experimental workflow for H_2_O_2_ treatment and digestion of NIH 3T3 cells expressing HA-tagged KRAS^G12C^. DTT: dithiothreitol, NEM: N-Ethylmaleimide, PRM: Parallel Reaction Monitoring (mass spectrometry analysis method). Created with Biorender.com. **(C)** Quantification of KRAS^G12C^ redox state in lysates from NIH 3T3 cells treated with control vehicle (PBS) or H_2_O_2_ (100 and 1000 μM). Representative extracted ion chromatograms of the KRAS C12-containing tryptic peptide labeled with NEM (representing reduced protein), d5-NEM (representing reversibly oxidized protein), or oxidized irreversibly to sulfinic acid (-SO_2_H). Peak area quantification based on mass spectrometry PRM analysis shows the contribution of dominant fragment ions.

With the observation that recombinant KRAS^G12C^ protein is rapidly oxidized *in vitro* upon addition of H_2_O_2_ (**Fig. S5**), we asked whether KRAS^G12C^ can be similarly oxidized in a cellular context. To test this hypothesis, we established NIH 3T3 cells stably expressing HA-tagged KRAS^G12C^. Following treatment with 0, 100 or 1000 μM H_2_O_2_ to induce KRAS oxidation, cells were harvested in a lysis buffer containing N-ethylmaleimide (NEM) to block reduced thiols and prevent artifactual oxidation. KRAS was immunoprecipitated, isolated via gel extraction and treated with DTT and then N-ethyl-d_5_-maleimide (d_5_-NEM) prior to trypsinization and targeted analysis of the G12C containing peptide by mass spectrometry (**Fig. 5B**). The analysis summarized in **Figure 5C** indicates that under basal conditions, KRAS^G12C^ resides in a mixture of reduced (NEM-labeled) and reversibly oxidized (d_5_-NEM-labeled) states, and that H_2_O_2_ addition shifts most of the protein to -SO_2_H, a redox state resistant to inhibitor binding. Interestingly, this modification is structurally very similar to aspartic acid, a prevalent oncogenic mutant (KRAS^G12D^) in human tumors (1,22).

### KRAS^G12C^ glutathionylated at C12 exhibits modestly altered biochemical properties

We and others have previously shown that KRAS and other RAS subfamily GTPases readily react with oxidizing agents like glutathione disulfide *in vitro* (14,15,23). In fact, oxidation of select RHO GTPases within the phosphoryl binding loop regions has been shown to alter GTPase function (14). With reversible oxidation observed for KRAS^G12C^ in NIH 3T3 cells, which could in part reflect glutathionylated protein, we hypothesized that KRAS^G12C^ glutathionylation may alter KRAS biochemical properties. First, we demonstrated that KRAS^G12C^ can indeed undergo stoichiometric glutathionylation *in vitro*. We found that modification of KRAS^G12C^ by oxidized glutathione occurred rapidly and to >90% modification. These studies were performed with the C118S mutant and the modification at the G12C cysteine was further confirmed by its interference with ABD-F modification (**Fig. 6A**).

**Figure 6.**
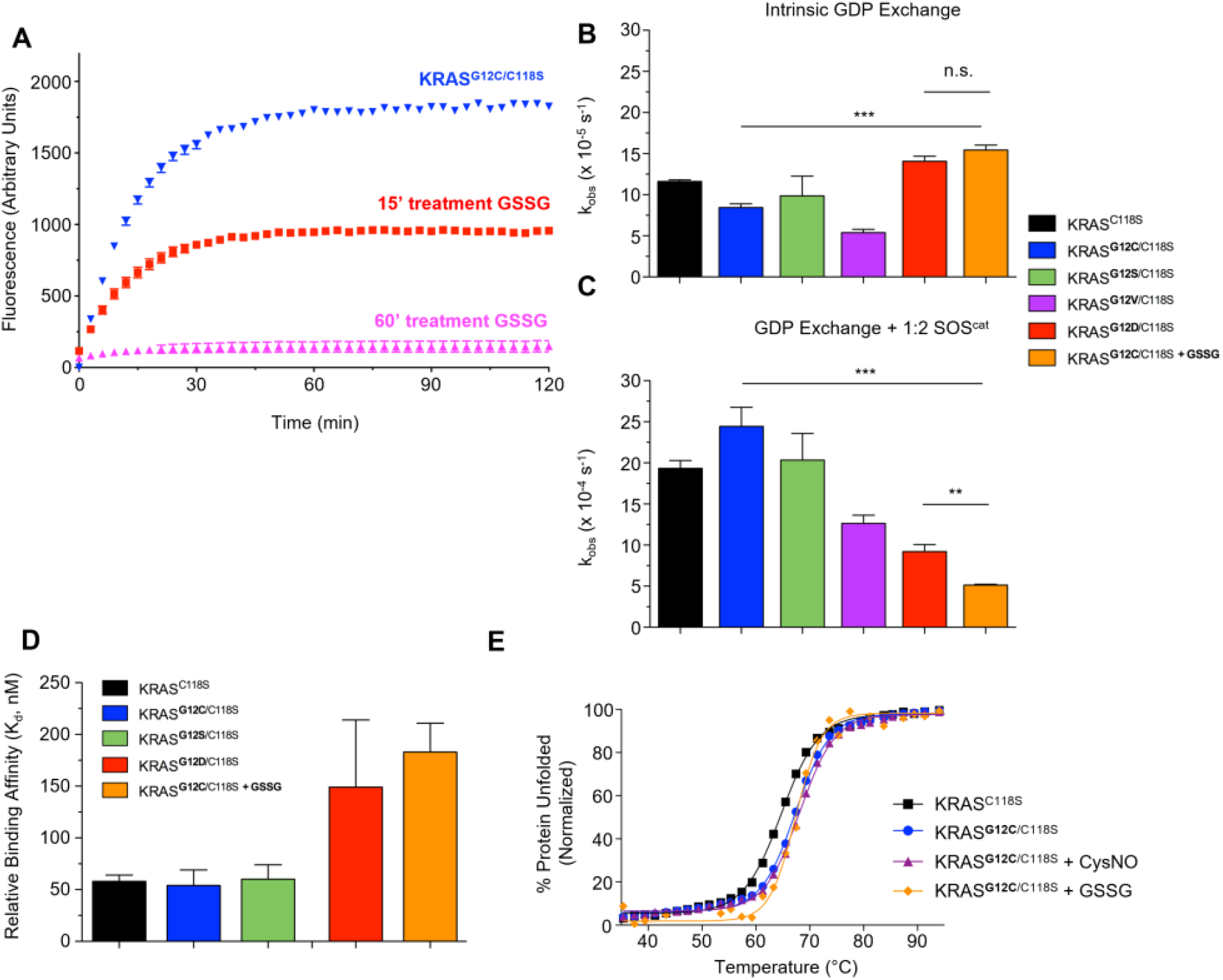
Glutathionylated KRAS^G12C^ shows altered biochemical function. **(A)** Relative ABD-F reactivity of KRAS^G12C/C118S^ pre-treated with oxidized glutathione (GSSG) for the indicated times at a 1:10 ratio. Following pre-treatment, KRAS (10 μM) was buffer exchanged and reacted with ABD-F (1 mM) at pH 8.0. No ABD-F reactivity as monitored by fluorescence increase is seen with KRAS^G12C/C118S^ after 1 h pretreatment with GSSG indicating complete modification of the reactive cysteine by GSSG. Error bars, mean ± s.e.m. **(B)** Intrinsic nucleotide dissociation rates of wild-type, mutant, and glutathionylated KRAS^G12C^. Nucleotide dissociation was measured by monitoring the decrease in ^mant^GDP fluorescence over time after addition of unlabeled GDP. Data shown are averaged from three or more independent experiments. ***, p ≤ 0.001, by one-way ANOVA. Error bars, mean ± s.e.m. **(C)** GEF-mediated nucleotide dissociation rates of wild-type, mutant, and glutathionylated KRAS^G12C^ measured as in panel B, but in the presence of the catalytic domain of the RAS GEF SOS1 (SOS^cat^) at a 1:2 ratio. Data shown are averaged from three or more independent experiments. ***, P ≤ 0.001; **, P ≤ 0.01, by one-way ANOVA. Error bars, mean ± s.e.m. **(D)** Dissociation constants (K_d_) of wild-type, mutant, and glutathionylated KRAS^G12C^ in complex with BRAF-RBD as determined by inhibition of nucleotide dissociation using a non-hydrolyzable analogue of fluorescent GTP, ^mant^GMPPNP. Data shown are averaged from three or more independent experiments. Error bars, mean ± s.e.m. **(E)** Circular dichroism melting temperatures of KRAS^G12C^ modified by CysNO (nitrosation) and GSSG (glutathiolation) for 60 min before analysis. Data shown are representative from three or more independent experiments.

While glutathionylation is a common reversible redox modification at cysteine residues, it is not the only form of oxidation that can occur; our data in **Fig. 5** and **Fig. S5** demonstrate oxidation of the G12C cysteine to sulfenic and sulfinic acid forms. As noted above, cysteine sulfinic acid (-SO_2_H) bears a striking resemblance to aspartic acid (-CO_2_H), suggesting that oncogenic G12D could behave similarly to this or other oxidized forms of KRAS^G12C^. Therefore, we also utilized other mutants at G12 including KRAS^G12D^ as a comparison to reduced and glutathionylated KRAS^G12C^. We hypothesized that substitutions at G12 of KRAS may alter nucleotide binding as the G12 residue interacts directly with GDP-GTP in native KRAS. To understand the impact of glutathionylation on KRAS^G12C^ nucleotide exchange activity, we pre-loaded glutathionylated protein (KRAS^G12C^ [-SSG]) with a fluorescent nucleotide analog (^mant^GDP), then added unlabeled GDP and monitored the decrease in fluorescence as a function of time to determine the rate of nucleotide dissociation. The intrinsic rate of nucleotide dissociation for KRAS^G12C^ (-SSG) was modestly elevated compared with unmodified KRAS^G12C^, and interestingly similar to KRAS^G12D^ (**Fig. 6B**).

As nucleotide exchange for KRAS is intrinsically slow without stimulation by RAS guanine nucleotide exchange factors (RAS GEFs), we tested the effect of the catalytic domain of the RAS GEF Son-of-Sevenless (SOS1) on KRAS-mutant nucleotide exchange rates. SOS1-stimulated GDP dissociation for KRAS^G12C^ (-SSG) was found to be 3-4-fold slower than unmodified KRAS^G12C^, indicating modestly impaired binding to SOS1; the slower exchange rate of the glutathionylated protein is similar to, but even slower than, those observed for KRAS^G12D^ (**Fig. 6C**). As the SOS1 interaction interface with KRAS involves the switch regions, we hypothesized that similar decreases in binding affinity would be observed for other RAS effectors, as well. In agreement with this, we observed that KRAS^G12C^ (-SSG) displayed ~3-fold decreased binding affinity to the RAS-binding domain (RBD) of BRAF as compared to unmodified KRAS^G12C^, yet similar affinity to that of KRAS^G12D^ (**Fig. 6D, Table S3**). These findings suggest that glutathionylated KRAS^G12C^, similar to KRAS^G12D^, can participate in signaling events through its interactions with regulatory proteins and effectors.

Lastly, we performed thermal denaturation analysis by CD to assess whether oxidative modification would significantly affect KRAS^G12C^ protein stability (**Fig. 6E**). We found that modification by glutathione or nitrosation from CysSNO treatment did not appreciably change KRAS melting temperature or cooperativity of unfolding (**Fig. 6E** and **Table S4**), suggesting that these oxidative modifications do not significantly affect protein stability.

#### MD simulations of KRAS^G12C^ and KRAS^G12D^ proteins

To better understand the structural implications of KRAS^G12C^ oxidation, we performed MD simulations to interrogate KRAS^G12C^ (-SO_2_^-^) and KRAS^G12C^ (-SSG) structure and dynamics for comparison with unmodified KRAS^G12C^ as well as KRAS^G12D^. The MD simulations were initialized from crystal structures of GDP-bound KRAS^G12C^ (PDB: 4LDJ) (24) and KRAS^G12D^ (PDB: 5US4) (25). For these comparisons, the G12C thiol was configured in protonated (-SH) or unprotonated states (-S^-^), modified by oxidation to sulfinate (unprotonated sulfinic acid, -SO_2_^-^) or glutathionylated (-SSG). From the backbone root-mean-square-deviation (RMSD) plots, we observed that all simulations equilibrated within the simulation time of 800 ns (**Fig. 7A** and **Fig. S6**). While only minor structural differences were observed among KRAS^G12C^ (-SSG), KRAS^G12C^ (-SH), KRAS^G12C^ (-S^-^) and KRAS^G12D^ systems (**Fig. 7A and Fig. S6A**), with an average RMSD between KRAS^G12C^ (-SSG) and KRAS^G12D^ trajectories of approximately 1.5 Å, differences in Switch dynamics were notable (Fig. **S6B**). Mapping the root-mean-square-fluctuation (RMSF) changes onto the three-dimensional structure of KRAS (**Fig. 7B**) shows that the oxidation of KRAS^G12C^ to the sulfinate form diminished Switch I dynamics relative to unprotonated (thiolate) KRAS^G12C^ (**Fig. 7B and 7D**) and glutathionylated KRAS^G12C^ (**Fig. S6B and S6D**). However, both KRAS^G12C^ (-SO_2_^-^) and KRAS^G12D^ exhibit residue-specific fluctuations and Switch dynamics on a similar scale. Another indication that the structure within the nucleotide binding region of the oxidized KRAS^G12C^ (-SO_2_^-^) is not significantly perturbed is the preservation of the full complement of GDP interactions observed in KRAS^G12D^ and unmodified KRAS^G12C^ (**Fig. 7C**). Furthermore, molecular docking of the known KRAS^G12D^ inhibitor MRTX-1133 to a representative, highly populated conformation extracted from the structural ensemble of KRAS^G12C^ (-SO_2_^-^) across the trajectory indicates strong binding in the P2-pocket. The ligand orientation and residue-specific interactions of MRTX-1133 with KRAS^G12C^ (-SO_2_^-^) are identical to the known X-ray structure of the MRTX-1133 bound to KRAS^G12D^ (PDB: 7RPZ) (26) (RMSD = 0.63 Å), which indicates the potential for KRAS^G12D^ inhibitors to target this oxidized form of KRAS^G12C^ **(Fig. 7E)**. Thus, the modeling data suggests that irreversible oxidation of KRAS^G12C^ to sulfinic acid yields a structure similar to KRAS^G12D^.

**Figure 7.**
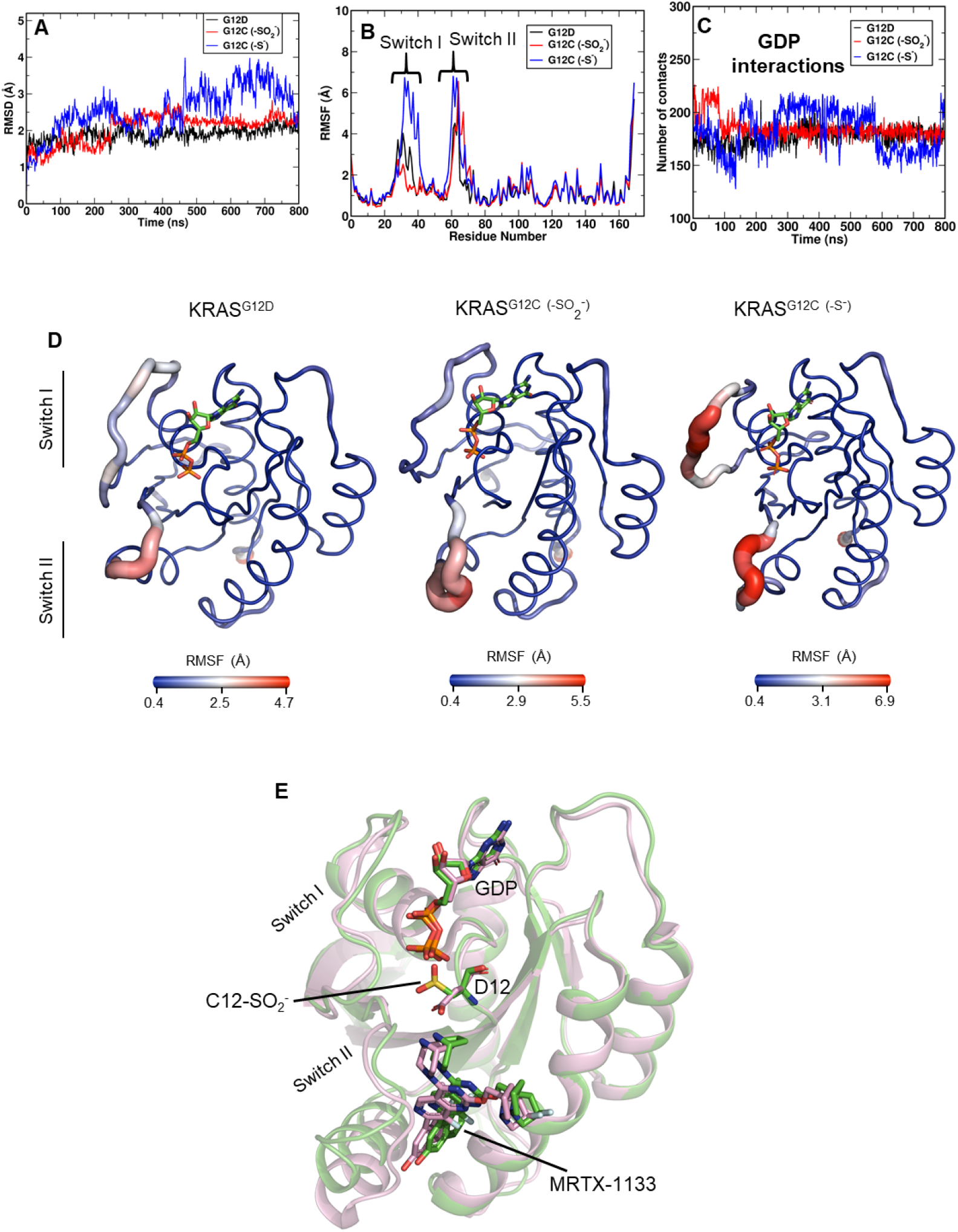
KRAS^G12C^ (-SO_2_^-^) and KRAS^G12D^ exhibit similar structural ensembles. Shown are **(A)** RMSD and **(B)** RMSF plots for GDP-bound KRAS^G12D^, KRAS^G12C^ (-SO_2_^-^) and KRAS^G12C^ (-S^-^). **(C)** Quantification of atomic interactions for RAS-bound GDP with surrounding residues in KRAS^G12D^, KRAS^G12C^ (-SO_2_^-^) and KRAS^G12C^ (-S^-^) indicate very similar GDP binding interactions in all three. **(D)** Sausage representation of KRAS^G12D^, KRAS^G12C^ (-SO_2_^-^) and KRAS^G12C^ (-S^-^) structures were extracted from respective MD trajectories and indicate that KRAS^G12D^ and KRAS^G12C^ (-SO_2_^-^) exhibit fluctuations in the KRAS Switch II region, whereas KRAS^G12C^ (-S^-^) exhibits fluctuations in both Switch I and Switch II. **(E)** Molecular docking of MRTX-1133, a known G12D inhibitor, to one of the highly populated structural ensembles of KRAS^G12C^ (-SO_2_^-^) (green) shows similar binding to KRAS^G12D^ (pink) as determined by X-ray crystallography (PDB: 7RPZ). The RMSD between MRTX-1133 bound KRAS^G12D^ and KRAS^G12C^ (-SO_2_^-^) is approximately 0.65 Å.

## DISCUSSION

Inhibition of mutant KRAS remains a highly pursued goal in drug discovery efforts for the treatment of human cancers. The discovery and application of KRAS^G12C^-specific inhibitors has been an exciting development in recent years, with several compounds achieving robust clinical success (8,9). However, treatment-acquired resistance remains a significant hurdle for targeted inhibition strategies. In part, a better understanding of how the reactivity of KRAS^G12C^ plays a role in inhibition kinetics will guide and improve further development of future inhibitors for this RAS mutant and others. To this end, our studies provide much-needed elucidation of the reactivity of the additional KRAS^G12C^ cysteine residue, C12, toward covalent inhibitors and oxidants. We developed an assay to inform the design and characterization of future KRAS inhibitors using the Y137W mutation, which was selected based on the location of a naturally-occurring Trp in some RAS family members. This substitution, which did not affect nucleotide exchange or protein stability, allowed for direct monitoring of rapid fluorescence changes associated with compound binding and ligation. Using this approach, we found that ARS-853 binds rapidly (only detectable at 5°C) with a K_d_ of 36 μM and is insensitive to pH from pH 7 – 8.5. However, the ligation reaction itself is pH dependent with a pK_a_ associated with k_inact_/K_i_ of 8.2 at both 5 and 20°C. Interestingly, assessment of C12 pK_a_ in the absence of inhibitor yielded a lower pK_a_ of 7.6 by both NMR and ABD-F reactivity approaches. Additionally, we found that KRAS^G12C^ is readily modified by cellular oxidants such as hydrogen peroxide and glutathione disulfide resulting in modifications that can alter KRAS structure and dynamics, with small reductions in regulatory and effector interactions. Notably, oxidation blocked covalent attachment of alkylating inhibitors.

With respect to the kinetic studies, we found the rate constants at 5°C to be consistent with a mechanism involving a pH-independent reversible binding step followed by pH-dependent irreversible covalent attachment of the inhibitor to C12. At 20°C, the binding step becomes too fast to monitor (complete in <1.5 ms) and determination of K_i_ relies only on data reflecting the hyperbolic dependence of the rate of the ligation step on the concentration of inhibitor. In our experiments at this temperature, only k_inact_/K_i_ can be reliably tracked given the high K_i_. When comparing our results with a previous study (10) (Hansen *et al.*) which included 1 mM DTT in the reaction buffer there were notable differences in the hyperbolic profiles at higher inhibitor concentrations (**Fig. S4**). The overlay of the datasets from the Hansen *et al*. and our MHT buffers at pH 7.5 shows that the steady increase in reaction rate (k_inact_) drops off at inhibitor concentrations >150 μM in the DTT-containing buffer but not in the MHT buffer, suggesting some rate limiting effect of the DTT-containing buffer at the higher inhibitor concentrations. While the previous study did not test inhibitor concentrations in these high ranges due to inhibitor insolubility (10), we did not encounter a problem with inhibitor solubility up to 300 μM. This could be due to the inclusion in our buffer of 3% DMSO, which did not affect the protein based on the kinetic assays, but likely improved inhibitor solubility. Our overall conclusions from the data at pH 7.5 and 20°C are that the K_i_ for ARS-853 is indeed high, at ~140 μM in the DTT-containing buffer of the previous study, or above 500 μM in our MHT buffer (**Fig. S4**). While Hansen *et al*. reported a k_inact_/K_i_ of 250 ± 90 M^-1^ s^-1^ (10) at pH 7.5, our findings estimate this at 510 ± 75 M^-1^ s^-1^ in the DTT-containing buffer used in the previous studies and 336 ± 45 M^-1^ s^-1^ in the MHT buffer. Note, however, that our findings show that the k_inact_/K_i_ value is highly sensitive to pH; at pH 7.5, it is only about 7% of the plateau value of 4,650 ± 190 M^-1^ s^-1^ determined for the fit to data in **Fig. 3E**. Thus, the main conclusions from our work and the previously published study of Hansen *et al*. (10) are that at 20°C and pH 7.5, the K_i_ for ARS-853 is quite dependent on the buffer components, but ranges from ~140 μM in the presence of DTT to much higher values that are difficult to determine and which are quite distinct from the nM IC50 values in cells (9). The k_inact_/K_i_, however, is also relatively high, at 250 M^-1^ s^-1^ (10) or up to 510 M^-1^ s^-1^ measured here at 20°C and pH 7.5.

Our studies have yielded some very interesting observations regarding the pK_a_ of the G12C cysteine, which in the absence of inhibitor is 7.6 ± 0.4 by two distinct approaches. The pK_a_ was first determined by tracking Cys reactivity toward an electrophile, ABD-F (**Fig. 1A & B**). The second approach employed direct monitoring of the C12 NH proton chemical shift across a pH range using NMR, which queries native protein and avoids addition of small molecules (**Fig. 1C & D**). These approaches indicate a shift of approximately one pH unit lower than an unperturbed Cys thiol group, enhancing the proportion of the reactive Cys thiolate form relative to the thiol at physiologically relevant pH values. This is an important attribute of KRAS^G12C^ that supports its reactivity toward electrophilic inhibitors as well as oxidants. Our findings contrast with the earlier study which used iodoacetamide and a discontinuous mass spectrometry-based approach to track the unmodified C12-containing peptide, where the pK_a_ was reported to be 9.0 ± 0.2 (10). While the origin of the differences in conclusions about pK_a_ are not clear, we feel that our two independent methods, and particularly the use of NMR, provide confidence that the pK_a_ of 7.6 for unliganded KRAS^G12C^ is an accurate result.

With our kinetic studies, we were also able to determine a functional pK_a_ extracted from the pH dependence of k_inact_/K_i_ for ARS-853 reaction (**Fig. 3E**). This value, at 8.21 ± 0.09 at 20°C, is higher than the ligand-free pK_a_ of 7.6 noted above, and also distinct from the value of 9.2 obtained previously using N-benzylacrylamide (10). While we again cannot clearly rationalize the discrepancies between our findings and those of the earlier study, we note that Hansen *et al*. data were collected at pH values well above pH 10, whereas in our hands the KRAS^G12C^ protein is unstable at such high pH values (our data were collected at pH values of 9.4 or lower). Taking a close look at the Michael addition mechanism and structural context of the C12 reaction with acrylamide-containing inhibitors, a plausible explanation for the different pK_a_ values for the inhibitor-free KRAS^G12C^ and its reactivity with inhibitor emerges. There has been considerable focus on a nearby lysine residue, K16, as critical to the activation of the acrylamide warhead, stabilizing the transition state during chemical ligation (10,11). As supported by detailed computational analyses, the protonated ammonium of K16 interacts with the carbonyl oxygen adjacent to the reactive vinyl group of the acrylamide, activating the terminal carbon for attack by the C12 sulfur (11). The reaction is then completed by protonation of the second carbon of the vinyl group from a Mg^2+^-bound water molecule. The C12 residue is therefore in a significantly different structural and electrostatic environment during reaction with acrylamide inhibitors than it is when the acrylamide is absent. Indeed, it is likely that the proximity of K16 to C12 in the ligand-free enzyme influences the C12 pK_a_ as it would be expected to stabilize the thiolate form, lowering the pK_a_. This may be the best explanation for our finding that the pK_a_ of the ligand-free enzyme is 7.6, whereas the pK_a_ observed for the reactivity of this Cys residue with the inhibitor is 8.2.

Another important point raised by our studies is that the protonation status of C12 does not affect binding of inhibitor as assessed for the fast step of ARS-853 reaction at 5°C (**Fig. 3B**). In fact, there is an indication that high pH may modestly decrease (rather than increase) the “on rate” (k_+1_) given the lower value and higher K_d_ at pH 9 (**Fig. 3B** and **Fig. S2**). This would be consistent with a role for K16 protonation in enhancing binding of the inhibitor to the enzyme in addition to its role in the chemical mechanism. As K16 is an essential player in the reaction mechanism, this hypothesis would be difficult to test experimentally.

Our data also support the oxidation sensitivity of the acquired C12, a factor which has been largely overlooked in the literature. We observed that two of the oxidized species likely to be found in KRAS^G12C^, sulfinic acid (which is irreversible) and glutathione connected through a mixed disulfide (which is reversible), are quite distinct in their effects on the protein structure and dynamics in the two switch regions. That this cysteine modification may serve as a mimic of the carboxylate sidechain of Asp (or *vice versa*) is well known and has been a useful tool in crystallographic studies (27). Indeed, our modeling and MD studies are supportive of the similarity between the sulfinate form of KRAS^G12C^ and KRAS^G12D^; in the face of oxidation blocking the covalent attachment of inhibitor, use of KRAS^G12D^ inhibitors may provide an additional approach through which the sulfinate-containing form could be inhibited (**Fig. 7** and **Fig. S6**). We also demonstrated herein the presence of reversibly oxidized C12 of HA-tagged KRAS^G12C^ in cells in the presence or absence of exogenous H_2_O_2_, as well as generation of the irreversibly-oxidized sulfinic acid form of C12 which quantitatively increases with addition of H_2_O_2_. While the nature of specific oxidative modifications at C12 in cells and tumors will be the subject of future studies, we note that it is well established that cancer cells generate high amounts of reactive oxygen species (ROS) and are more likely to exhibit higher levels of oxidative protein modifications (28,29). Moreover, RAS signaling itself promotes ROS generation, so redox regulation is highly relevant for KRAS-driven tumor growth and treatment (30,31).

Finally, our studies have shown that a clinically efficacious inhibitor, AMG 510 (also known as Sotorasib and trademarked as Lumakras™), also exhibited very rapid equilibration of binding, although in this case the binding and dissociation rate constants were so high at 5°C that binding equilibration was complete within the dead time of the stopped flow instrument (**Fig. 4A**). Further, while the K_i_ for AMG 510 was above the highest inhibitor concentration used (300 μM) and higher than that for ARS-853 (**Fig. S3**), the k_inact_/K_i_ second order rate constant for ligation was 5 - 9 fold higher for AMG 510 than for ARS-853 (**Table S1**). Expansion of our technologies to assess groups of acrylamide-based inhibitors with varying linkers and scaffolds and comparisons with known potency of such inhibitors with KRAS-driven cancer cells in culture or during patient treatments is likely to provide much-needed insight into mechanistic reasons for variations in potency. In addition, assessment of KRAS^G12C^ oxidation status in tumors prior to treatment may offer new opportunities for tailoring treatments to the patients most likely to respond to this class of covalent inhibitors.

## MATERIALS AND METHODS

### Protein purification

Truncated human KRAS4B (residues 2–169) was expressed from a modified pET21 bacterial expression vector containing an N-terminal 6x-His purification tag followed by a Tobacco Etch Virus (TEV) protease cleavage site. SOS^cat^ (residues 564–1049, pPROEX HTb) and BRAF-RBD (residues 149-232, pET28a) contained a similar vector architecture. All 6xHis-tagged proteins were expressed in Rosetta2 (DE3) cells and purified following the Qiagen Nickel NTA purification protocol and the 6xHis tags were removed using TEV protease. For pGEX vectors, proteins were purified following the Glutathione Sepharose™ 4B purification protocol (Amersham Pharmacia Biotech). The GST-tag was cleaved overnight using thrombin protease while dialyzing in wash buffer. If necessary, the proteins were further purified by size exclusion chromatography (Superdex-75 10/300 GL column; GE Life Sciences) and judged greater than 95% pure by SDS-PAGE analysis.

### ABD-F modification assays

KRAS was exchanged into ABD-F modification buffer (15 mM HEPES, 15 mM MES, 5 mM MgCl_2_, 50 mM NaCl, 200 μM DTPA) supplemented with 10 mM DTT and allowed to incubate for 30 m on ice. At the same time, ABD-F modification buffer without DTT was sparged with N2 gas to remove dissolved oxygen. The protein was exchanged into this buffer to remove DTT from the sample prior to reaction with ABD-F.

For modification with ABD-F, 20 μM KRAS was added in 100 μL of reaction buffer with a pH that was pre-determined to a black 96-well plate. To a separate plate, ABD-F was added to 2 mM in 100 μL of reaction buffer at the same pH as the corresponding well in the original plate.

Using a multi-channel pipette, ABD-F was added to the KRAS plate to start the reaction. The fluorescence of the reaction was monitored using a SpectraMax M5 plate reader over a pH range of 5.8 to 8.5. The excitation wavelength for ABD-F is 389 nm, and the emission wavelength is 513 nm. The mutant KRAS^G12C/C118S^ was used for pK_a_ analyses.

To generate oxidized KRAS^G12C^ modified by CysNO (13) or GSSG (14), reduced KRAS^G12C/C118S^ was reacted with 100 molar equivalents of either oxidizing agent for one hour in a buffer supplemented with 100-fold excess of GDP at 37°C. Following incubation, KRAS was buffer exchanged to remove oxidizing agents and complete modification was verified using ABD-F analyses.

### NMR analysis

Purification of ^13^C, ^15^N-enriched KRAS^G12C/C118S^ required no modifications to the purification protocol described above. NMR spectra were acquired at 25°C on a Bruker Avance 850 NMR spectrometer (19.97 T field strength) using a cryogenic (TCI) 5 mm triple-resonance probe equipped with z-axis gradient. Two-dimensional (2D) ^1^H-^15^N HSQC experiments were performed as previously described (32). For NMR analysis, ^13^C, ^15^N-enriched KRAS proteins were exchanged into a buffer containing 20 mM Tris-Maleate, 40 mM NaCl, 5 mM MgCl_2_ and 20 μM GDP, supplemented with 5% D2O. The NMR data were processed using Topspin (v3.6.1, Bruker) and the spectra were visualized using SPARKY (33). Backbone resonance assignments of KRAS^G12C^ bound to GDP were previously obtained (18,19).

### Circular dichroism (CD) spectroscopy

KRAS was exchanged into a buffer containing 10 mM KH_2_PO_4_/K_2_HPO_4_ at pH 7.4 and diluted to 15 μM. Generation of oxidized KRAS^G12C^ modified by CysNO (13) or GSSG (14) was performed as described earlier. MgCl_2_ and GDP were added to a final concentration of 500 and 80 μM respectively before analysis. Experiments were performed on a Jasco J-815 CD Spectrometer, with samples measured using a 1-mm cuvette. A thermal melt scan from 20° to 90°C was performed to determine the melting temperature (Tm) at which half of the protein is unfolded.

### Guanine nucleotide exchange and protein binding assays

Loading of KRAS proteins with fluorescent nucleotide analogs was performed as previously described (34). For loading of the non-hydrolyzable nucleotide analog ^mant^GMPPNP (mGMPPNP, Jena Biosciences), KRAS protein was incubated with alkaline phosphatase beads (Sigma Aldrich) and 20-fold excess nucleotide overnight with gentle rotation at 4°C. Alkaline phosphatase and excess nucleotide were removed with a buffer exchange using a PD-10 desalting column. Purified protein was checked for nucleotide loading by HPLC to confirm >90% binding of the respective nucleotide (35). In brief, nucleotide dissociation was initiated by addition of 1000-fold excess of unlabeled GDP at 25°C. The rate of dissociation was monitored by the change in fluorescence at an excitation wavelength of 365 nm and emission at 435 nm, using a SpectraMax M5 plate reader with a 384-well Greiner plate. Fluorescent nucleotide dissociation curves were fit to a single exponential decay equation using GraphPad Prism. Loading of KRAS with ^mant^GDP (mGDP, Jena Biosciences) was performed following previously published methods (34). Nucleotide exchange assays were performed using a Cary Eclipse Fluorescence Spectrophotometer (Agilent), as previously described (34). The minimal catalytic fragment of the RASGEF SOS1 (SOS^cat^) was used to stimulate nucleotide dissociation along with the addition of 1000-fold excess of unlabeled nucleotide. All experiments were performed in triplicate.

Quantitative binding of KRAS to RAS effector domains was performed as previously described (36). In brief, KRAS loaded with mGMPPNP (1.5 μM) was incubated with increasing concentrations of BRAF-RBD (20 mM HEPES, 50 mM NaCl and 5 mM MgCl_2_ at pH 7.4). Nucleotide dissociation was initiated by addition of 1000-fold excess of unlabeled nucleotide at 25°C. The rate-of-dissociation was monitored by the change in fluorescence at an excitation wavelength of 365 nm and emission at 435 nm, using a SpectraMax M5 plate reader with a 384-well Greiner plate. Fluorescent nucleotide dissociation curves were fit to a one-phase exponential decay equation using GraphPad Prism. Extrapolated nucleotide dissociation rates were fit against the effector concentration using previously published methods (37). All experiments were performed in triplicate.

### Stopped-flow kinetic studies

KRAS and varying concentrations of inhibitor (ARS-853, AMG 510) were prepared in “MHT” buffer containing 50 mM MES, 50 mM HEPES, 50 mM Tris, 50 mM NaCl, 5 mM MgCl_2_, 3% (v/v) DMSO and titrated with NaOH over a pH range from 6.5 to 9.4. The KRAS was first reduced by adding 10 mM DTT and incubating at 20°C for 30 minutes, and then passed over a PD-10 desalting column to remove excess DTT, then the protein was exchanged into a buffer containing 10 mM HEPES at pH 7.5, with 50 mM NaCl, 5 mM MgCl_2_, and 10 μM GDP. Samples were loaded into the syringes of an Applied Photophysics SX.18MV stopped-flow spectrophotometer and allowed to equilibrate to 5 or 20°C. The fluorescence changes were recorded after sample mixing via monitoring at 90° relative to incident light with excitation at 280 nm and emission >320 nm. Separate time courses were used to investigate the fast initial changes (0.2-1 s) and the much slower phase (100-1000 s) of decreasing fluorescence signal. Data were analyzed using Applied Photophysics software and fit to single (or double) exponentials as appropriate. To obtain rate constants and inhibition parameters, data from all (four or more) inhibitor concentrations at a given pH and 5°C were fit globally using KinTek Global Kinetic Explorer (version 10.2.0) (20,38) using Equation 2 (both steps are taken to be reversible, but the second step returns a reverse rate that is essentially 0). At 20°C, where the fast phase of fluorescence change is too fast to detect, hyperbolic dependence of the reaction rate on inhibitor concentration ([I]) is expected and observed, but in many cases the data are collected at [I] around or below the apparent K_i_, thus the “Kst” method of Ken Johnson (20) was used, which returns relatively reliable k_inact_/K_i_ values and errors, even at very high K_i_. Briefly, data were fit to equation 3 using Kaleidagraph software, version 4.5.4:

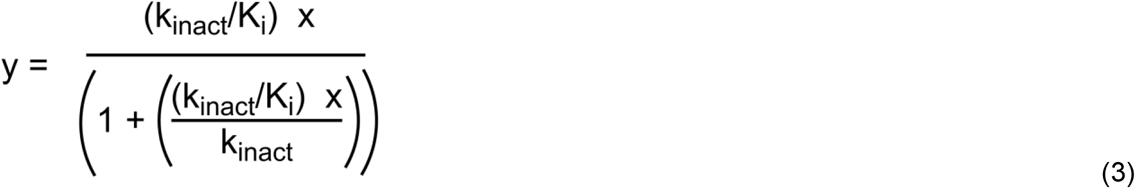

### Cell culture

NIH 3T3 cells were maintained in DMEM supplemented with 10% Colorado calf serum (CCS, Colorado Serum Company) in a humidified chamber with 5% CO_2_ at 37°C. Full-length human KRAS4B G12C was ectopically expressed from the retroviral expression vector pBABE containing an N-terminal HA-tag (Addgene #58901). Viral particles were generated by transient transfection of each expression vector into HEK 293T cells using Fugene6 (Promega) with the PCL-10A1 packaging system according to the manufacturer’s recommended protocol. Infection of cell lines was performed in growth medium supplemented with 8 mg/mL polybrene, with antibiotic selection beginning 48 h after transduction.

### Mass spectrometry analysis of KRAS^G12C^ modification

For detection of intracellular oxidation of KRAS^G12C^, cells transfected with HA-tagged KRAS^G12C^ were treated with 100 μM and 1 mM hydrogen peroxide for 15 min. Cells were harvested in the lysis buffer supplemented with 100 mM N-ethylmaleimide (NEM). HA-KRAS^G12C^ was immunoprecipitated using the Pierce HA-Tag IP Kit (26180, Thermo Scientific, Waltham, MA) and separated by SDS-PAGE. The gel band containing the HA-KRAS was excised, treated with 10 mM DTT at 56°C for 1 hour followed by 30 mM N-ethyl-d_5_-maleimide (d_5_-NEM) for 30 min at room temperature and subjected to in-gel tryptic digestion overnight at 37°C (10 ng/mL trypsin in 50 mM NH_4_HCO_3_ were freshly made to cover the gel pieces). Peptides were extracted by 50% ACN / 5% formic acid, and dried using a SpeedVac (SPD1010, Thermo Scientific, Waltham, MA). The peptides were purified using C18 spin tips (Cat. No. 87784, Thermo Scientific, Waltham, MA) and solubilized in water containing 5% acetonitrile (ACN) / 0.1% formic acid for LC-MS/MS analysis.

Samples were analyzed on a LC-MS/MS system consisting of an Orbitrap Velos Pro Mass Spectrometer (Thermo Scientific, Waltham, MA) and a Dionex Ultimate3000 nano-LC system (Thermo Scientific, Waltham, MA). An Acclaim PepMap 100 (C18, 5 μm, 100 Å, 100 μm x 2 cm) trap column and an Acclaim PepMap RSLC (C18, 2 μm, 100 Å, 75 μm x 50 cm) analytical column were employed for peptide separation. The mobile phases were 5% ACN in water (solvent A) and 80% ACN in water (solvent B) both of which contained 0.1% formic acid. Mass spectra were acquired in positive ion mode by alternating MS1 and MS2 scans with targeting 563.8181 m/z and 566.3335 m/z for identification of NEM and d_5_-NEM labeled LVVVGA<u>C</u>GVGK peptide, respectively, and 517.2892 m/z to identify sulfinylated LVVVGACGVGK peptide. Skyline (MacCoss Lab Software, University of Washington, Seattle, WA) was used for target peptides detection, peak feature extraction, and peak area calculation for quantitative data analysis. Peak areas were normalized using the total ion current (TIC) which is the sum of all peaks in the chromatogram acquired by complementary MS1 scan event. NIST mass spectral library was utilized to confirm peak selection for the analysis.

For mass spectrometry analysis of the reactions between reduced and oxidized Cys-light KRAS^G12C^ (KRAS^G12C/CL^) with ARS-853, purified KRAS protein was reduced with addition of 10 mM DTT for 30 min at 25°C. The samples were buffer exchanged using a P6 Bio-Gel column that was pre-equilibrated with 50 mM ammonium bicarbonate (ABC). Protein concentration was determined from the solution absorbance at 280 nm (ε = 23930 M^-1^ cm^-1^). Oxidation was achieved by addition of 1-10 molar equivalents of H_2_O_2_ for 10 min at 25°C with 250 RPM shaking using a thermomixer. The reaction was quenched by passing through a P6 Bio-Gel spin column pre-equilibrated with ABC. Formation of KRAS^G12C/CL^ oxidation products was monitored by ESI-TOF mass spectrometry. Reduced or oxidized KRAS^G12C/CL^ was mixed with 1.5 molar equivalents of ARS-853 in ABC buffer for 5 min at 25°C with 250 RPM shaking using a thermomixer. The reaction was quenched, and formation of KRAS^G12C/CL^ adducts with ARS-853 were monitored by Electrospray Ionization Time-of-Flight mass spectrometry (ESI-TOF MS).

Analysis of intact KRAS^G12C^ proteins was performed on an Agilent 6120 MSD-TOF system operating in positive ion mode with the following settings: capillary voltage of 3.5 kV, nebulizer gas pressure of 30 psi, drying gas flow of 5 L/min, fragmentor voltage of 175 V, skimmer voltage of 65 V, and drying gas temperature of 325°C. Samples were introduced via direct infusion at a flow rate of 20 μL/min using a syringe pump. Mass spectra were acquired over the range of 600-3200 m/z and then averaged and deconvoluted, and ion abundance was quantified using Agilent MassHunter Workstation software vB.02.00. Relative ion abundances were used to determine the reaction progress.

### Molecular dynamics (MD) simulations

The crystal structures of GDP-bound KRAS^G12C^ (PDB accession code 4LDJ) (24) and GDP-bound KRAS^G12D^ (PDB accession code 5US4) (25) were used as starting structures for simulations. For simulations of KRAS^G12C/Y137W^, the Y137W substitution was introduced in the KRAS^G12C^ X-structure (PDB: 4LDJ). MD simulations of GDP-bound KRAS^G12C^ were performed with C12 in the protonated (-SH), thiolate (-S^-^), sulfinate (-SO_2_^-^) and glutathionylated (-SSG) states. All missing residues and atoms were modelled using Modeller-9v18 tool prior to MD simulations (39). To generate the model of KRAS^G12C^ oxidized to the sulfinate (-SO_2_^-^), starting with the KRAS^G12C^ structure, we replaced C12 with an Asp residue, then replaced the β carbon of the Asp sidechain with a sulfur atom. We used the CHARMM36 topology (40,41) of the Asp residue as a basis to generate C12 (-SO_2_^-^) forcefield parameters. To generate the KRAS conformation with glutathionylated G12C (-SSG), initially the backbone atoms of Cys22 from human glutaredoxin (PDB accession code 1B4Q) (42) were aligned to the G12C backbone from KRAS^G12C^. Subsequently, the glutathione (Glu-Cys-Gly) tripeptide was extracted from glutaredoxin and attached to the side-chain of G12C through S-S disulfide linkage. The modeled KRAS^G12C^ (-SSG) structure was energy minimized to remove atomic clashes between the G12C (-SSG) modification and the surrounding residues prior to MD simulations. The topology of the G12C (-SSG) moiety was generated by following step-by-step instructions from http://www.ks.uiuc.edu/Training/Tutorials/science/topology/topology-tutorial.pdf before modification to match the GROMACS convention. The starting structure was immersed in a periodic water box and the system charge was neutralized by adding an appropriate number of Na^+^ counterions. Each system was optimized using 10,000 steps of steepest-descents energy minimization. Subsequently, position restrained MD simulations were performed for 1 ns in isothermal–isobaric ensemble (constant temperature and constant pressure) by restraining backbone heavy atoms of protein and GDP. Production simulations were performed for 1000 ns with a simulation time step of 2 fs. The temperature and pressure were maintained at 310K and 1 bar by employing V-rescale thermostat (43) and Parrinello-Rahman barostat (44) respectively. Electrostatic interactions were estimated using particle mesh Ewald method (45) with cutoff distance at 1.2 nm. The van der Waals interactions were terminated at cut-off value of 1.2 nm and LINCS algorithm (46) was used to constrain all bonds with H-atoms. All simulations were performed using GROMACS-2020 software package (47) and CHARMM36 forcefield (40,41). Trajectory analyses was performed using GROMACS built-in tools and inhouse scripts. All structural figures were rendered using PyMOL visualization software (48).

Molecular docking of MRTX-1133 to the representative conformation extracted from highly populated structural ensemble of KRAS^G12C^ (-SO_2_^-^) was performed using the Hex docking program (49). Hex employs real orthogonal spherical polar basis functions to represent surface shape and charge distributions of protein and ligand molecules, which is subsequently used to estimate probable docked complex conformations through FFT calculations. Hex evaluates docking score between receptor and ligand as a function of the six degrees of translational and rotational freedom in a rigid body docking search.

## Funding

This research was supported in part by the following grants: R35 GM135179 to L.B.P. R35 GM134962 to S.L.C., and U01 CA215848 to C.M.F. The authors wish to acknowledge also the Wake Forest Baptist Comprehensive Cancer Center Proteomics and Metabolomics Shared Resource, as well as the Crystallography and Computational Biosciences Shared Resource, supported by the National Cancer Institute’s Cancer Center Support Grant award number P30 CA012197.

## Conflict of interest

The authors declare that they have no conflicts of interest with the contents of this article.

## Notes

### Competing Interest Statement

The authors have declared no competing interest.

