## Supporting Information for "Kinetic and Redox Characterization of KRAS G12C Inhibition"

List of Material Included

<u>Figures</u>

**Figure S1. Y137W mutation does not perturb the structural dynamics of KRAS$^{G12C}$.**

**Figure S2. Stopped-flow kinetic studies of KRAS$^{CCLW}$ and ARS-853 at 5 °C.**

**Figure S3. Additional comparison of stopped-flow kinetics of KRAS$^{CCLW}$ with ARS-853 vs. AMG 510.**

**Figure S4. Comparison of stopped-flow kinetics data for KRAS$^{CCLW}$ with ARS-853 with MHT buffer versus the buffer from Hansen et al., 2018.**

**Figure S5. KRAS$^{G12C}$ is rapidly oxidized after 10 min treatment with $H_2O_2$.**

**Figure S6. Glutathionylated KRAS$^{G12C}$ and KRAS$^{G12C}$ thiolate show altered switch dynamics.**

<u>Tables</u>

**Table S1. Summary of $k_{inact}/K_i$ values obtained for inhibitors at 20 °C and 3 pH values.**

**Table S2. Kinetic parameters at 20 °C and pH 7.5 of KRAS$^{CCLW}$ with ARS-853 compared with previously reported results (1).**

**Table S3. Binding affinity data for KRAS mutants as shown in Figure 6D.**

**Table S4. Thermal melt data for KRAS mutants as shown in Figure 6E.**

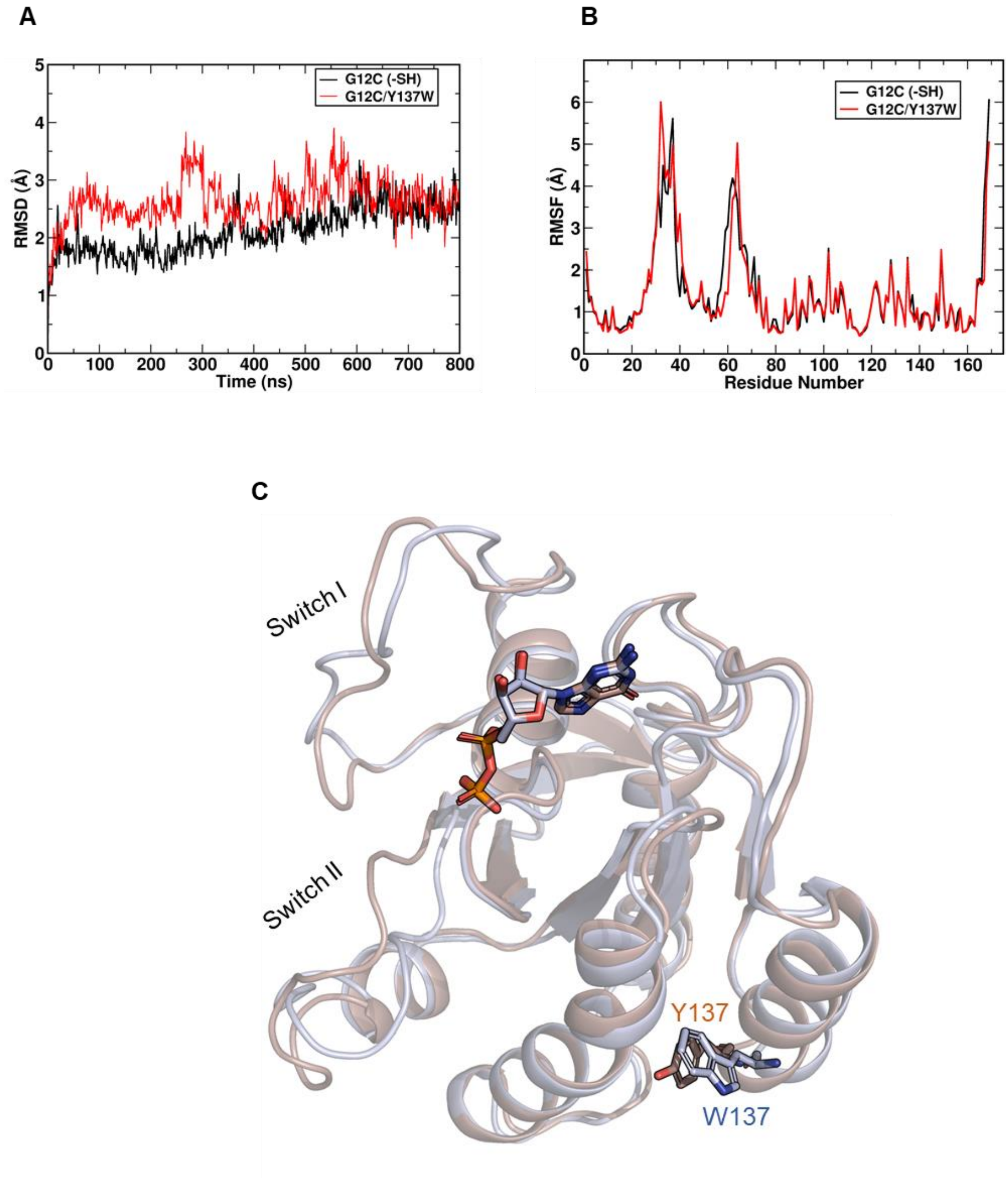


**Figure S1. The KRAS$^{G12C/Y137W}$ ~~RAS$^{Y137W}$~~ variant does not perturb the structure and dynamics of KRAS$^{G12C}$.** (**A** and **B**) RMSD and RMSF plots obtained from MD trajectories of KRAS$^{G12C}$ and KRAS$^{G12C/Y137W}$ indicate that the Y137W substitution does not significantly alter KRAS$^{G12C}$ structure and dynamics. **(C)** Overlay of highly populated structural ensembles of KRAS$^{G12C}$ (~~gray~~brown) and KRAS$^{G12C+Y137W}$ (light blue) extracted from respective MD trajectories. Protein is shown in ribbon and GDP is shown in stick representation.

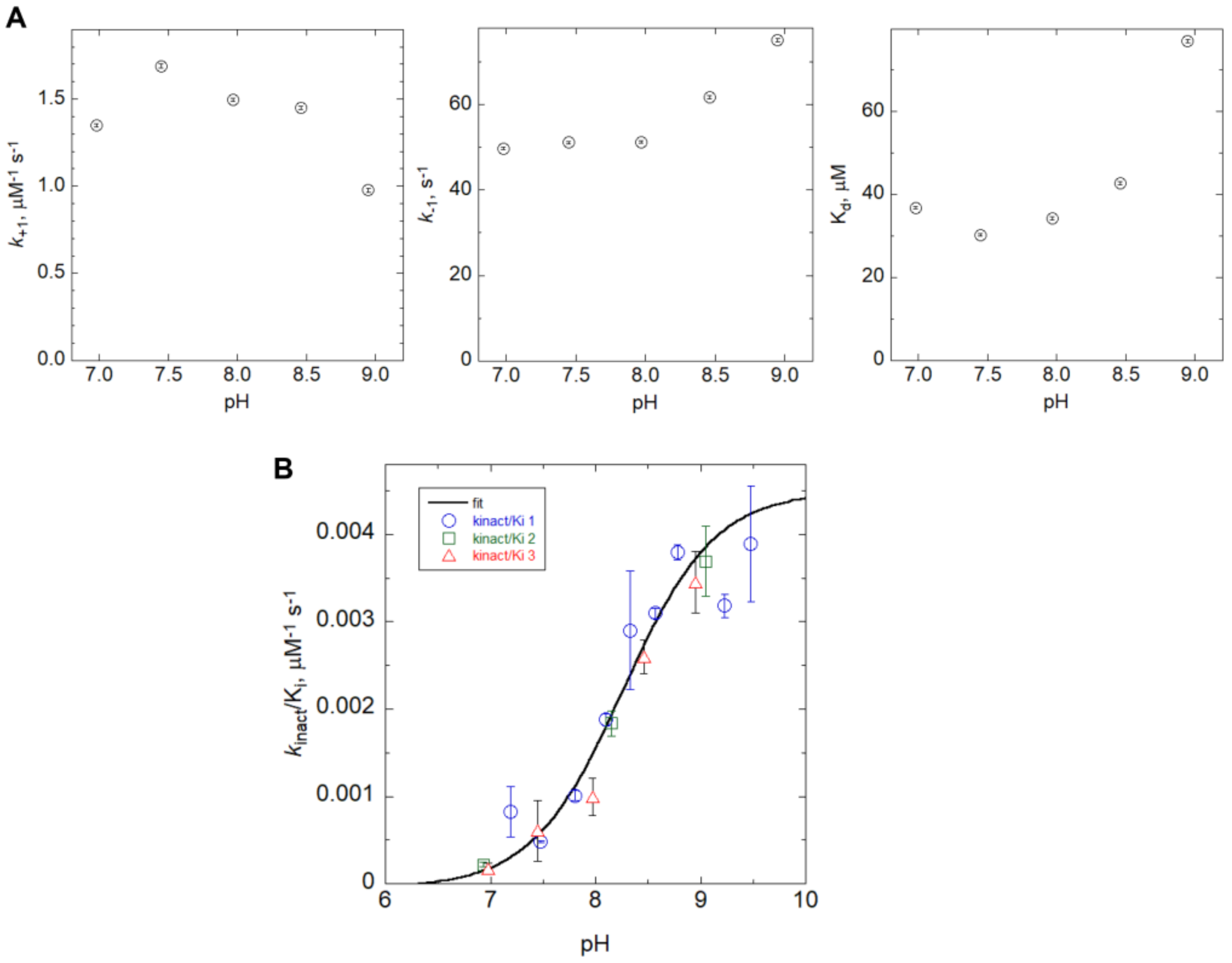


**Figure S2. Stopped-flow kinetic studies of KRAS$^{CCLW}$ and ARS-853 at 5 °C.** (**A**) Kinetic rate constants and $K_d$ for the fast step of KRAS$^{CCLW}$ and ARS-853 show little pH sensitivity at 5 °C. (**B**) Second order rate constants ($k_{inact}/K_i$) for the slow step of KRAS$^{CCLW}$ and ARS-853 show similar pH sensitivity at 5 °C as compared to 20 °C (Fig. 3E). Three colors and marker shapes represent three independent trials, all used in the final fit. Error bars, mean ± **s.e.m**.

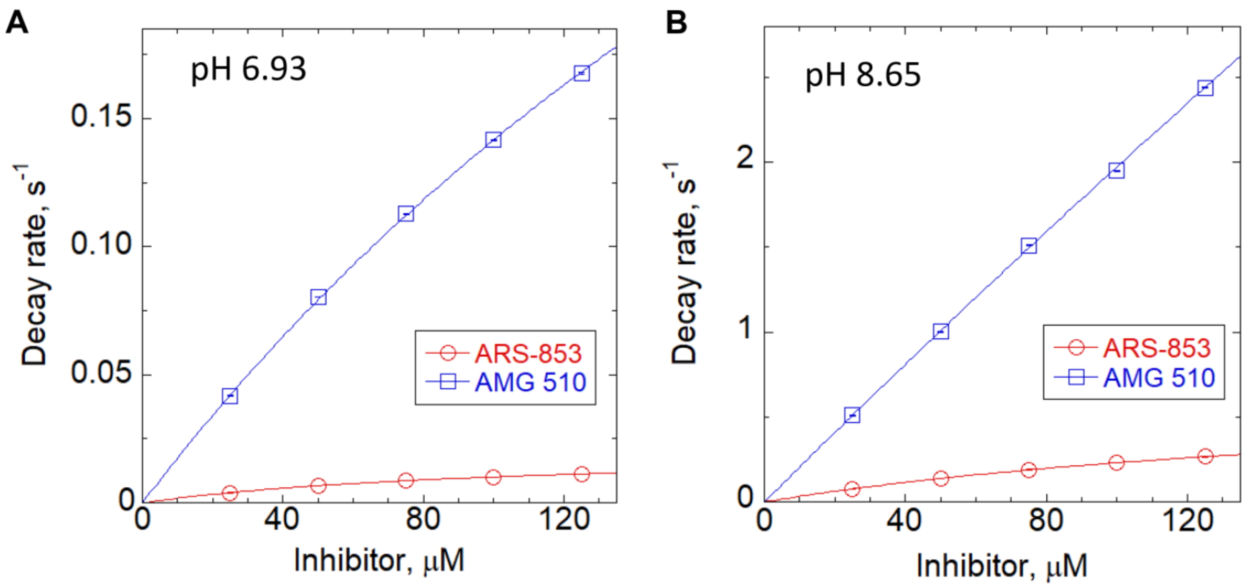


**Figure S3. Additional comparison of stopped-flow kinetics of KRAS$^{CCLW}$ with ARS-853 vs. AMG 510. (A and B)** Plots of the pseudo first order rate constants versus inhibitor concentrations of KRAS$^{CCLW}$ with AMG 510 versus ARS-853 at pH 6.93 (**A**) and pH 8.65 (**B**) at 20°C, showing fits to the "Ksp" treatment of the hyperbolic Michaelis-Menten equation as described in methods (poorly determined $K_i$ values are similar to or higher than the highest inhibitor concentrations used). Reactions were monitored in a MES-HEPES-Tris buffer.

**Table S1. Summary of $k_{inact}/K_i$ values obtained for inhibitors at 20 °C and 3 pH values.**

| | pH 6.93 | | pH 8.61 (in Fig. 4) | | pH 8.65 | |
|---|---|---|---|---|---|---|
| | AMG 510 | ARS-853 | AMG 510 | ARS-853 | AMG 510 | ARS-853 |
| $k_{inact}/K_i$ ($M^{-1}$ $s^{-1}$) | 1790 (± 10) | 200 (± 5) | 14000 (± 100) | 2770 (± 10) | 20600 (± 100) | 3450 (± 10) |
| $k_{inact}/K_i$ (AMG vs. ARS) | 9.0-fold higher | | 5.0-fold higher | | 6.0-fold higher | |

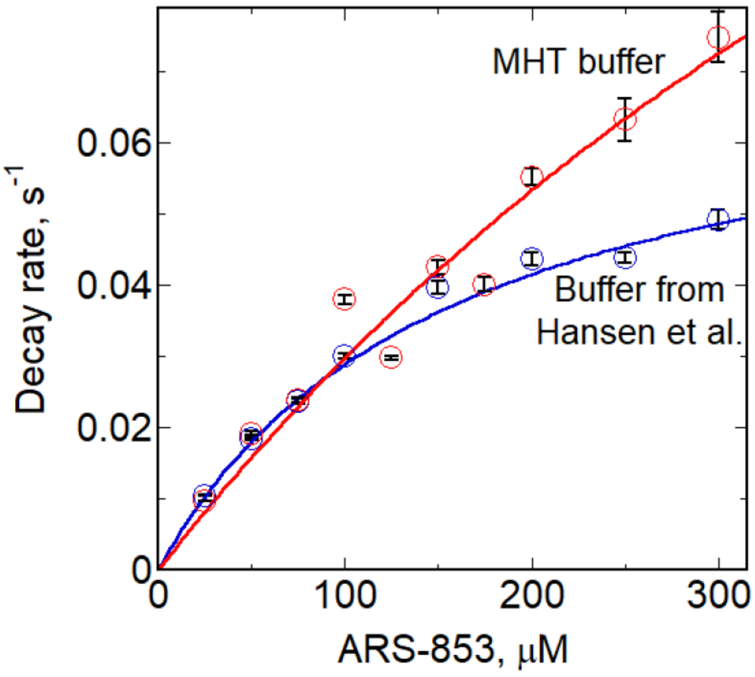


**Figure S4. Comparison of stopped-flow kinetics data for KRAS$^{CCLW}$ with ARS-853 with MHT buffer versus the buffer from Hansen et al., 2018 (1).** Plots of the pseudo first order rate constants versus inhibitor concentrations of KRAS$^{CCLW}$ with ARS-853 in a MES-HEPES-Tris (MHT) buffer (red), or the buffer composition used by Hansen et al. containing DTT (blue) at 20°C.

**Table S2. Kinetic parameters at 20 °C and pH 7.5 of KRAS$^{CCLW}$ with ARS-853 compared with previously reported results (1).**

| Buffer: | MHT buffer[a] | Hansen et al. buffer[b] | Literature data[c] |
|---|---|---|---|
| $K_i$ (μM) | >500 | 142 ± 19 | 200 ± 90 |
| $k_{inact}$ ($s^{-1}$) | >0.15 | 0.072 ± 0.004 | 0.05 ± 0.023 |
| $k_{inact}/K_i$ ($M^{-1}$ $s^{-1}$) | 336 ± 45 | 510 ± 75 | 250 ± 20 |

[a]Data were fit using the "Ksp" approach as described in Methods.

[b]Data were fit to equation 1 (in the main text).

[c]Data reported by Hansen et al. (1).

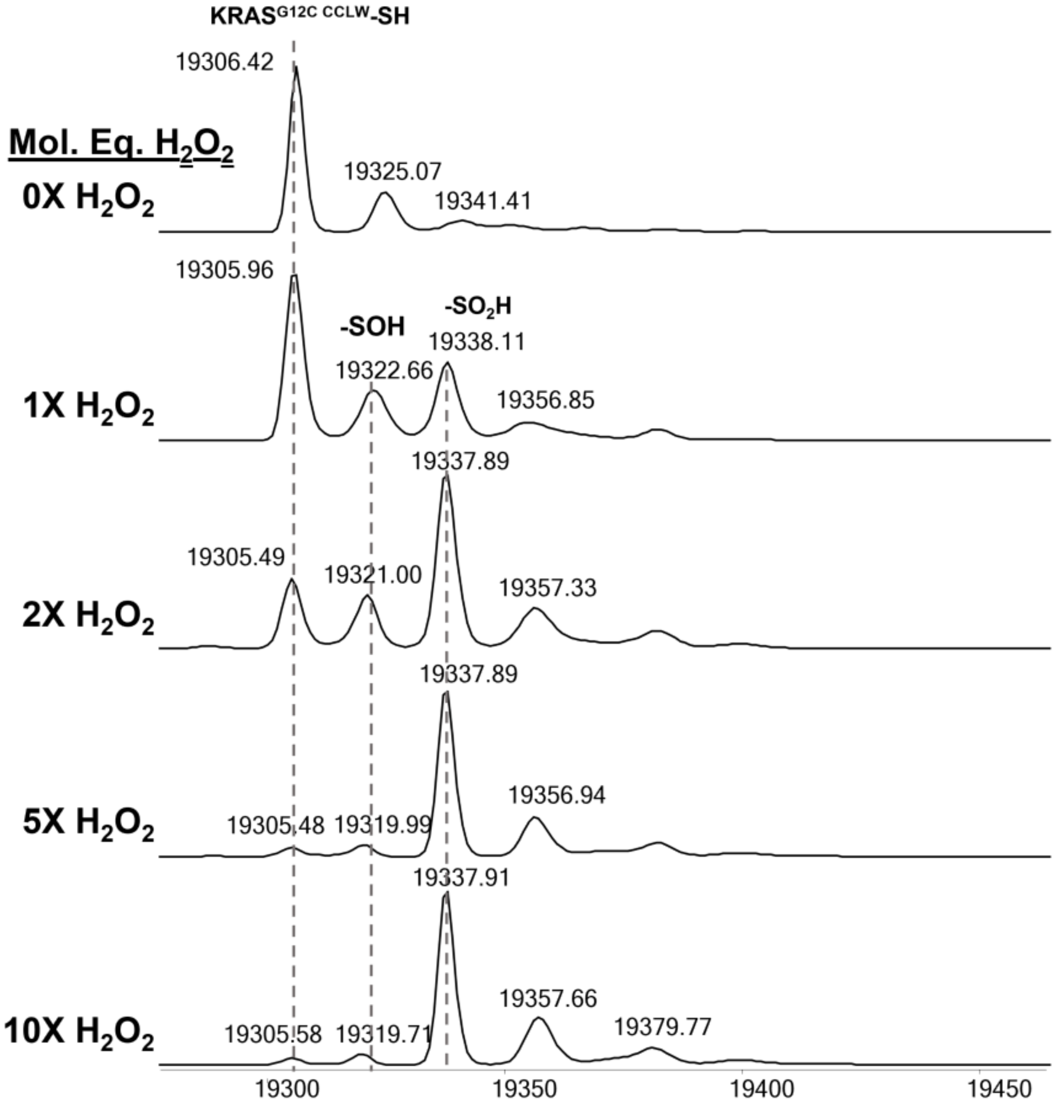


**Figure S5. KRAS$^{G12C}$ is rapidly oxidized after 10 min treatment with $H_2O_2$.** Increasing molar equivalents of $H_2O_2$ show increased formation of sulfenic (-SOH) and sulfinic acid (-$SO_2$H) modifications.

**Table S3. Binding affinity data for KRAS mutants as shown in Figure 6D.[a]**

| | $K_d$ (nM) |
|---|---|
| $KRAS^{C118S}$ | 58 ± 6 |
| $KRAS^{G12C/C118S}$ | 54 ± 15 |
| $KRAS^{G12C/C118S}$ + GSSG | 183 ± 28 |
| $KRAS^{G12D/C118S}$ | 149 ± 65 |
| $KRAS^{G12S/C118S}$ | 60 ± 14 |

[a] Calculated binding affinities of wild-type and mutant KRAS proteins in complex to BRAF-RBD as determined by inhibition of nucleotide dissociation. Data shown are averaged from three or more independent experiments.

**Table S4. Thermal melt data for KRAS mutants as shown in Figure 6E.[a]**

| | $T_m$ (°C) |
|---|---|
| $KRAS^{C118S}$ | 64.5 ± 1.5 |
| $KRAS^{G12C/C118S}$ | 67.1 ± 1.1 |
| $KRAS^{G12C/C118S}$ + CysNO | 67.8 ± 2.5 |
| $KRAS^{G12C/C118S}$ + GSSG | 67.0 ± 2.8 |

[a] Calculated circular dichroism melting temperatures of KRAS proteins. Data shown are averaged from three or more independent experiments.

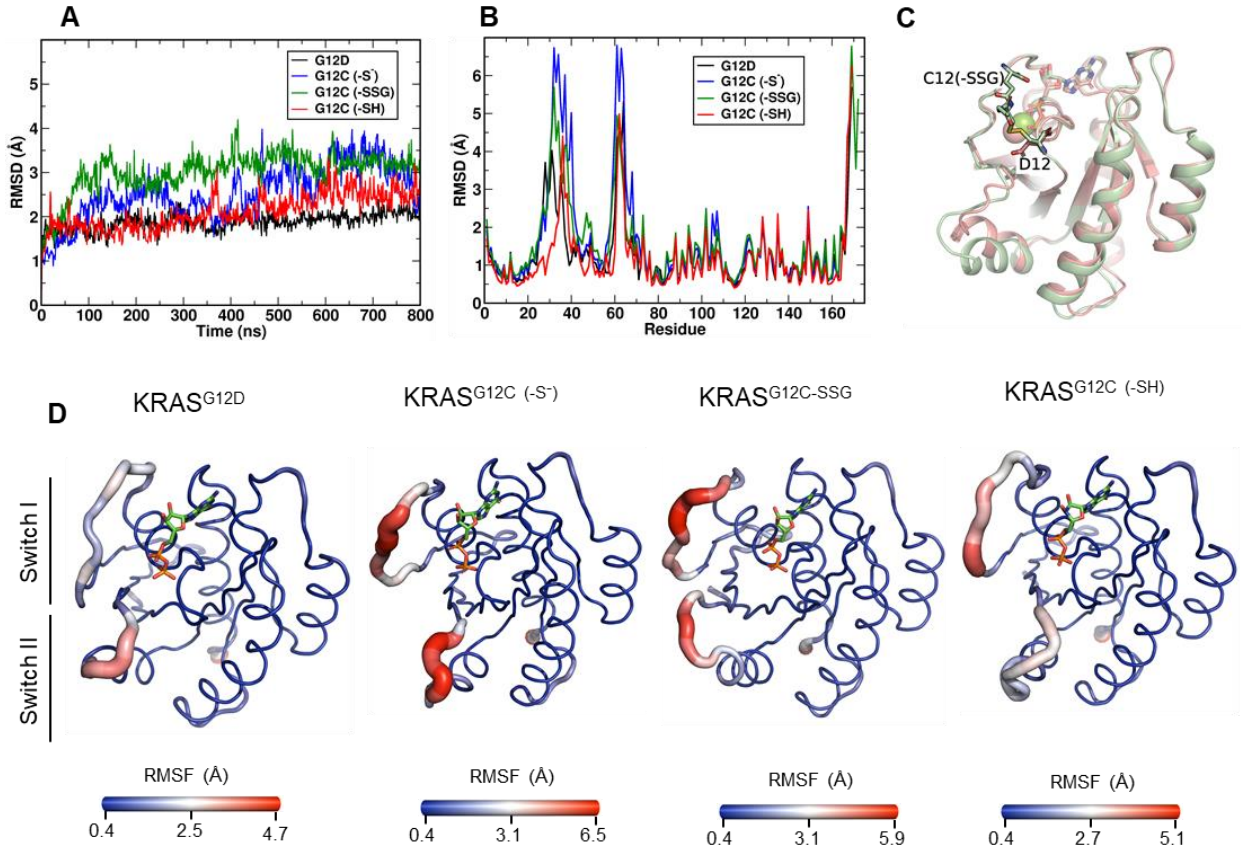


**Figure S6. Glutathionylated KRAS$^{G12C}$ and KRAS$^{G12C}$ thiolate show altered switch dynamics.** (**A** and **B**) Molecular dynamics (MD) trajectories of GDP-bound KRAS$^{G12D}$, as well as KRAS$^{G12C}$ in the protonated (-SH), thiolate (-S$^-$), and glutathionylated state (-SSG). KRAS$^{G12C}$ (-SSG) show similar structures. **(C)** Ribbon diagram overlay of modeled KRAS$^{G12C}$ (-SSG) (salmon) and KRAS$^{G12D}$ (green). G12C-SSG and G12D side-chains are represented as sticks. **(D)** Sausage representation of KRAS$^{G12D}$, KRAS$^{G12C}$ (-S$^-$), KRAS$^{G12C}$ (-SSG), and KRAS$^{G12C}$ structures extracted from respective MD trajectories. Glutathionylation of the KRAS$^{G12C}$ thiol and deprotonatation of KRAS$^{G12C}$ (thiolate) increase fluctuations in the KRAS Switch regions as compared to reduced, protonated KRAS$^{G12C}$ (-SH) and KRAS$^{G12D}$.

<u>Reference</u>


1. Hansen, R., Peters, U., Babbar, A., Chen, Y., Feng, J., Janes, M. R., Li, L. S., Ren, P., Liu, Y., and Zarrinkar, P. P. (2018) The reactivity-driven biochemical mechanism of covalent KRAS(G12C) inhibitors. *Nature structural & molecular biology* **25**, 454-462
